# Characterizing multiscale cellular remodeling through hierarchical deconvolution with HIDE-Deconv

**DOI:** 10.64898/2026.08.24.746754

**Authors:** Dennis Voelkl, Sarah Bolz, Austin Rayford, Thomas Sterr, Malte Mensching-Buhr, Nicole Seifert, Julia Arp, Jana Tauschke, Laurenz Engel, Cornelia Schuster, Thomas Stevenson, Helena U. Zacharias, Michael Altenbuchinger, Franziska Görtler

## Abstract

Disease-associated tissue remodeling involves coordinated changes in cellular populations across multiple levels of cellular resolution, from broad cell classes to specialized subpopulations. While single-cell RNA sequencing enables detailed characterization of cellular heterogeneity, bulk RNA sequencing remains far more widely available in large cohorts and clinical studies. However, most existing deconvolution approaches estimate cellular composition at only a single level of resolution and do not explicitly incorporate hierarchical relationships between cell populations.

Here, we present HIDE-Deconv, a hierarchical deconvolution framework that jointly estimates cellular compositions across multiple levels of a user-defined cell-type hierarchy. HIDE-Deconv combines reference construction, hierarchical deconvolution and downstream analytical workflows within a reproducible framework that can be applied to bulk RNA-seq data using single-cell reference atlases. By jointly modeling multiple hierarchy levels, the method produces cellular composition estimates that remain consistent across cellular resolutions while capturing both broad lineage-level and fine-grained cell-type information.

Using pseudobulk RNA-seq benchmark datasets generated from single-cell data, HIDE-Deconv achieved strong predictive performance while jointly modeling cellular compositions across multiple levels of a cell-type hierarchy. Application to lung adenocarcinoma revealed biologically and clinically distinct tumor microenvironment states that differed in cellular composition, transcriptional programs, response to PD-L1 blockade and overall survival. Analysis of independent peripheral blood mononuclear cell datasets from patients with sepsis, COVID-19 and systemic lupus erythematosus further demonstrated that disease-associated immune remodeling becomes apparent at different levels of cellular resolution depending on the underlying disease process.

Together, these results demonstrate that hierarchical deconvolution enables biologically relevant cellular remodeling to be investigated across multiple levels of cellular organization and provides a practical framework for extracting interpretable cellular information from bulk transcriptomic data.

**Author summary:** Bulk RNA sequencing remains widely used in large biomedical studies, but measurements represent mixtures of many different cell populations. Existing deconvolution methods typically estimate cellular composition at a single level of cellular resolution and do not account for hierarchical relationships between broad cell classes and their specialized subpopulations.

We developed HIDE-Deconv, a framework for hierarchical cell-type deconvolution from bulk RNA-seq data. HIDE-Deconv combines reference construction, deconvolution and downstream analyses within a reproducible workflow and estimates cellular composition across multiple levels of cellular resolution simultaneously.

Using both benchmark and disease datasets, we show that hierarchical modeling improves the consistency of cellular composition estimates and helps reveal biologically relevant remodeling patterns that become apparent at different levels of cellular resolution.

## 1 Introduction

Changes in cellular composition are a hallmark of many diseases, including cancer, infection and autoimmune disorders (1). In addition to transcriptional changes within individual cells, disease-associated processes alter the abundance of cellular populations, thereby reshaping tissue organization and function (2). Consequently, the cellular composition provides important biological and clinical information and can support biomarker discovery, patient stratification and the identification of molecular subtypes associated with therapeutic response and patient outcomes (3; 4).

Single-cell RNA sequencing (scRNA-seq) enables transcriptomic profiling at the cellular level and has substantially improved the characterization of complex tissues (5). Despite its impact on research, routine clinical application of scRNA-seq remains limited by experimental costs, throughput and technical biases affecting cell recovery (6; 7). Bulk RNA-seq, in contrast, remains widely employed owing to its scalability, established workflows and comparatively low cost. However, bulk RNA-seq measures aggregate gene expression across all cells within a sample and therefore does not directly resolve the underlying cellular composition. Computational deconvolution methods therefore aim to infer the cellular composition from bulk transcriptomic measurements (8).

A variety of deconvolution approaches have been proposed, including regression-based methods such as Digital Tissue Deconvolution (DTD), CIBERSORTx, Rectangle and EPIC (9; 10; 11; 12), probabilistic approaches such as BayesPrism (13) and deep-learning-based methods such as Scaden (14). While these approaches have substantially improved the cellular deconvolution of bulk transcriptomic data, accurate high-resolution cell-type deconvolution remains challenging (8; 15). Closely related cell-types often exhibit highly similar transcriptional profiles, whereas rare populations contribute only weakly to the overall expression signal. Consequently, cell-type annotations are often aggregated into broader cellular categories to improve robustness (16). While this can increase prediction accuracy, biologically relevant changes occurring within specific cellular subpopulations may be obscured. As a result, biological interpretation can depend strongly on the level of cellular resolution at which deconvolution is performed.

The hierarchical organization of cell-types (17; 18**?** ) provides a natural framework for addressing this trade-off between robustness and cellular resolution. Specialized cell types are organized into higher-level cellular lineages, forming multilevel structures that are reflected in established cell ontologies. Rather than treating each level of cellular resolution independently, hierarchical approaches can use information from higher-level cell populations to support the estimation of closely related and low-abundance cell-types. This can improve robustness while preserving the ability to investigate the cellular composition at different levels of resolution (16; 19).

In our previous work, we introduced Digital Tissue Deconvolution (DTD) (9) and Hierarchical Cell-Type Deconvolution (HIDE) (19). DTD addressed the difficulty of resolving rare and closely related cell populations by learning gene-specific weights that emphasize genes particularly informative for individual cell-types. Building on this concept, HIDE incorporates hierarchical cell-type relationships through a top-down deconvolution strategy that estimates broad cellular lineages before resolving increasingly fine-grained cell-types. By leveraging information across hierarchy levels, HIDE ensured hierarchically consistent cellular composition estimates across multiple levels of cellular resolution.

Other methods have also incorporated hierarchical or multiresolution cell-type annotations into the deconvolution process. For example, BayesPrism (13) and HiDecon (16) incorporate broad cellular lineages and finer cellular subtypes within hierarchical frameworks. Existing approaches differ substantially in how information across hierarchy levels is incorporated during training and inference. While methods such as HIDE perform deconvolution sequentially across different hierarchy levels, other approaches derive broader cellular abundances through aggregation of fine-grained predictions. Despite these advances, hierarchical information is typically incorporated only partially during model training or prediction. In addition, HIDE is based on a fixed three-level hierarchy and optimizes gene weights separately for each hierarchy level, limiting the integration of information across the hierarchy during model training.

Beyond these methodological challenges, practical barriers continue to limit the adoption of hierarchical deconvolution approaches. Deconvolution studies typically require multiple processing steps, including preprocessing of single-cell reference data, reference construction, model training, deconvolution, visualization and downstream statistical analyses. Because these steps are often distributed across multiple software tools, substantial technical expertise is frequently required to perform reproducible analyses. Although recent frameworks such as Deconomix (20) have demonstrated the benefits of integrating deconvolution-related functionality within a common software environment, comprehensive support for hierarchical deconvolution workflows remains limited.

To address both the methodological and practical limitations of existing hierarchical deconvolution approaches, we developed HIDE-Deconv. Methodologically, HIDE-Deconv replaces sequential hierarchy-aware inference with a joint optimization framework in which all hierarchy levels contribute simultaneously to a shared objective function. By allowing hierarchical information to directly influence model training, HIDE-Deconv supports deconvolution across hierarchies of varying depth and explicitly incorporates information from all hierarchy levels during model training.

HIDE-Deconv is distributed as an open-source Python package (https://github.com/dvoelkl/HIDE-deconv) that provides an integrated command-line interface and end-to-end workflows for hierarchical deconvolution. Starting from annotated single-cell reference data, bulk RNA-seq profiles and optional clinical metadata, the framework supports reference construction, model training, hierarchical deconvolution and downstream analyses within a unified workflow.

We evaluated HIDE-Deconv using synthetic bulk RNA-seq benchmark datasets generated from a lung adenocarcinoma (LUAD) single-cell atlas and compared its performance with established and recently developed deconvolution methods. In these benchmark analyses, HIDE-Deconv achieved a strong overall predictive performance compared to existing hierarchical approaches while maintaining consistency across hierarchy levels. We further applied the complete HIDE-Deconv workflow to a LUAD cohort treated with PD-L1 inhibitors and to peripheral blood mononuclear cell (PBMC) datasets from patients with sepsis, COVID-19 and systemic lupus erythematosus (SLE) to investigate disease-associated cellular remodeling across different levels of cellular resolution.

## 2 Materials and methods

### 2.1 Hierarchical deconvolution model

HIDE-Deconv extends the weighted least-squares deconvolution framework introduced in Digital Tissue Deconvolution (DTD). A detailed derivation of the original DTD objective function is provided in Supplementary Methods Section S1.1. Briefly, DTD estimates cellular compositions using a weighted least-squares formulation with learned gene-specific weights. HIDE-Deconv generalizes this framework to hierarchical cellular annotations by jointly incorporating multiple levels of cellular resolution during model training and inference.

Rather than optimizing each hierarchy level independently, HIDE-Deconv jointly incorporates information across the hierarchy. Cellular compositions are estimated at the finest level of cellular resolution, whereas abundances at coarser hierarchy levels are obtained through hierarchical projections. During training, hierarchy-specific gene weights are optimized jointly across all hierarchy levels, allowing both broad lineage-associated and fine-grained cell-type-specific information to contribute to model training.

To incorporate hierarchical cell-type annotations, we introduce a projection matrix *A_l_* ∈ ℝ*^ql×q^* where *q* denotes the number of cell types at the finest hierarchy level and *q_l_* the number of cell types at hierarchy level *l*. The projection matrix maps the finest-grained cellular composition *C* onto the corresponding cellular representation at hierarchy level *l*.

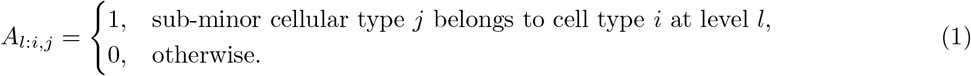

By construction, each *A_l_* projects compositions from the finest-grained cellular resolution onto the corresponding representation at hierarchy level *l*. We define *l* = 1 as the finest resolution level, for which *A*_1_ = *I_q×q_* is the identity matrix.

For each hierarchy level *l*, *X_l_* ∈ ℝ*^p×ql^* denotes the reference matrix containing the *q_l_* cell-type profiles defined at that resolution. Each reference matrix is weighted by a level-specific gene-weight matrix *G_l_* = diag(*g_l_*) ∈ ℝ*^p×p^*.

Using these definitions, the weighted least-squares problem of of the DTD algorithm 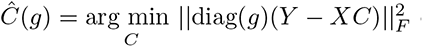 can be extended across all hierarchy levels as

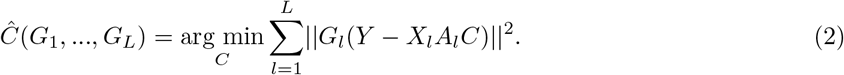

Defining

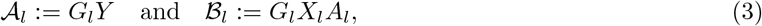

and stacking the resulting level-specific matrices, the optimization problem reduces to a standard least-squares problem,

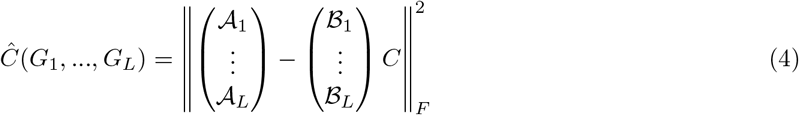

with a closed form solution for *C* with fixed *G_1_*,…*G_L_* given by

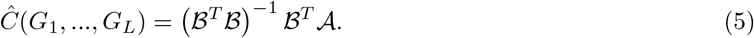

As with DTD, negative abundance estimates are set to zero after optimization, resulting in non-negative cellular compositions. The optimal gene weights *G_l_* are obtained by minimizing the outer loss function

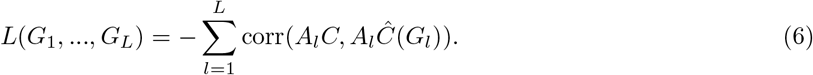

The loss function maximizes the Pearson correlation between the true and estimated cellular compositions across all hierarchy levels. Estimates at coarser levels of the hierarchy are obtained by projecting the predicted composition of the finest-resolution level using the corresponding projection matrices. After optimization, the learned gene-weight matrices *G*_1_*, . . . , G_L_* are fixed and applied to bulk RNA-seq samples. Cellular compositions are then estimated by solving the corresponding weighted least-squares problem and subsequently projected to coarser hierarchy levels using the matrices *A_l_*.

### 2.2 Datasets, feature selection and reference construction

Benchmark analyses were based on the extended LUAD single-cell atlas of Salcher et al. (21). After quality control, the dataset comprised 518,874 cells from 126 patients and 16,348 genes. To prevent information leakage, the contributing studies were split into independent training and test partitions for benchmark generation and model evaluation. Additional preprocessing and partitioning details are provided in Supplementary Methods Section S1.2.1.

For PBMC analyses, we used the Immune Health Atlas published by Gong et al. (22). After preprocessing, the reference dataset comprised 155,894 cells from 108 donors and 21,857 genes (additional details in Supp. Methods S1.2.2).

To evaluate the practical applicability of HIDE-Deconv, we analyzed bulk RNA-seq datasets from patients with LUAD, sepsis, COVID-19 and systemic lupus erythematosus (SLE). The LUAD analysis was based on 110 pre-treatment tumor samples from the SU2C-MARK cohort (23). PBMC analyses were performed using publicly available bulk RNA-seq datasets from patients with sepsis (GSE279448), COVID-19 (GSE152418) and SLE (GSE112087). A summary of the analyzed bulk RNA-seq datasets is provided in Supplementary Table S1. Reference profiles were constructed by averaging normalized gene expression across cells belonging to the same annotated cell-type population. Feature selection was performed on the finest level of the cell-type hierarchy by ranking genes according to their variance across reference profiles. The 5,000 highest-ranking genes were retained by default, although the number of selected genes can be specified by the user. The resulting feature set was used for model training and deconvolution analyses. For benchmark analyses, feature selection and reference construction were performed exclusively on the training partition to prevent information leakage between training and test data.

### 2.3 Cross-platform normalization between single-cell and bulk RNA-seq data

To account for systematic differences between single-cell and bulk RNA-seq platforms, we estimated gene-specific correction factors based on matched single-cell-derived pseudobulk and real bulk RNA-seq expression distributions. Correction factors were estimated independently from the samples subsequently used for deconvolution. The resulting gene-specific scaling factors were applied prior to deconvolution to improve cross-platform compatibility between reference and bulk expression profiles. A detailed description of the normalization procedure is provided in Supplementary Methods Section S1.3.

### 2.4 Benchmark design

We compared HIDE-Deconv against a diverse set of widely used reference-based deconvolution methods representing several methodological classes. The benchmark included weighted least-squares approaches (DTD (9) and HIDE (19)), probabilistic methods (BayesPrism (13)), reference-based regression methods (EPIC (12) and MuSiC (24)), partial-reference deconvolution (PREDE (25)), and the recently proposed Rectangle framework (11).

All methods were evaluated using the same study-level train-test split and benchmark pseudobulk datasets. Methods capable of operating at multiple levels of cellular resolution were evaluated at both major-and sub-minor-cell-type resolution. For methods lacking native hierarchical support, major-cell-type abundances were additionally obtained through aggregation of sub-minor predictions according to the cell-type hierarchy. Method-specific implementation details are summarized in Supplementary Table S1.4 and Supplementary Methods Section S1.4.1.

Benchmark datasets were generated from the independent LUAD test partition by aggregating expression profiles from randomly selected cells into pseudobulk samples with known cellular compositions. Ten independent benchmark datasets comprising 1,000 pseudobulk samples each were generated and used throughout all benchmarking analyses.

Predictive performance was assessed using Pearson correlation, Spearman correlation and normalized mean absolute error (NMAE). Hierarchical consistency was assessed by comparing directly estimated major-cell-type abundances with abundances obtained through hierarchical aggregation of sub-minor-cell-type predictions.

### 2.5 Application analyses

Deconvolved cellular compositions were used for downstream biological analyses, including dimensionality reduction, clustering, differential expression analysis and cancer hallmark enrichment analyses. Unless stated otherwise, all downstream analyses and visualizations, including PLS-DA, clustering, differential abundance analyses, differential gene expression analyses, survival analyses and heatmap visualizations, were performed using functionality provided by the HIDE-Deconv framework. Detailed descriptions of these procedures are provided in Supplementary Methods S1.5.

## 3 Results

### 3.1 HIDE-Deconv: a joint optimization model for hierarchical deconvolution

HIDE-Deconv combines hierarchical deconvolution, hierarchy-aware gene weighting and downstream analytical workflows within a unified framework (Figure 1). Unlike conventional deconvolution methods that operate at a single level of cellular resolution, HIDE-Deconv jointly models cellular compositions across multiple levels of a user-defined cell-type hierarchy while maintaining consistency between hierarchy levels. During training, information from all hierarchy levels contributes simultaneously to a shared optimization objective, enabling both broad lineage-associated and fine-grained cell-type-specific information to influence model training.

**Fig 1.**
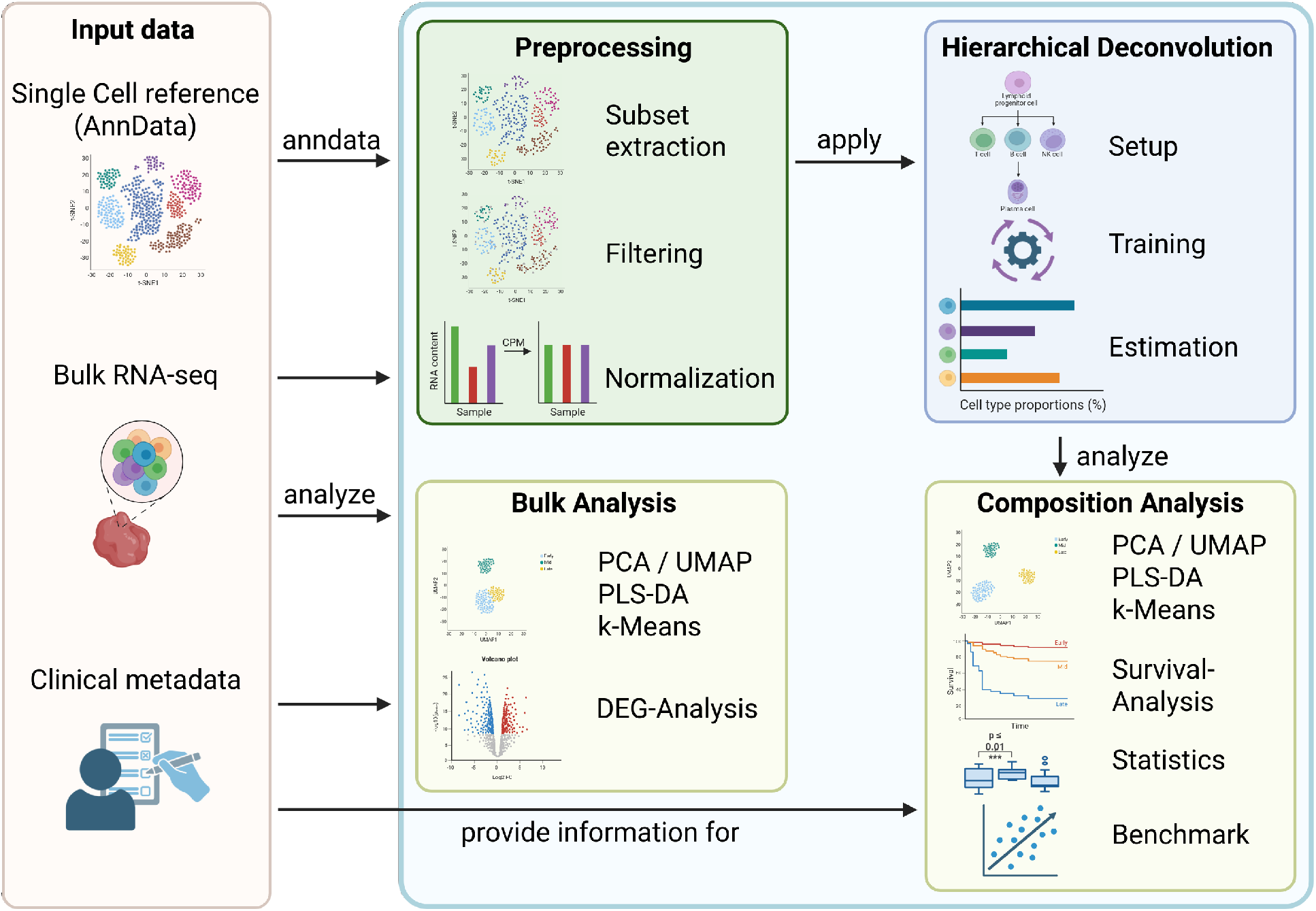
Overview of the HIDE-Deconv framework. HIDE-Deconv combines single-cell reference data, bulk RNA-seq data and optional clinical metadata within a unified workflow. The framework supports preprocessing, hierarchical deconvolution and downstream analyses including dimensionality reduction, clustering, differential expression, survival analysis, statistical testing and benchmarking. HIDE-Deconv can be accessed through a Python API or command-line interface.

Beyond the hierarchical model itself, HIDE-Deconv provides an end-to-end software framework supporting reference construction, model training, deconvolution and downstream analyses within a unified workflow. The framework can be applied to bulk RNA-seq datasets using annotated single-cell reference atlases and optional clinical metadata. Integrated analysis modules support dimensionality reduction, clustering, differential abundance testing, differential gene expression analysis, pathway analysis, survival analysis and benchmarking.

HIDE-Deconv is distributed as an open-source Python package and can be executed on local workstations, institutional computing clusters and secure computing environments. By operating entirely on local infrastructure, the framework can be applied to sensitive patient-derived datasets without requiring transfer of data to external servers or cloud services. The software is available through GitHub (https://github.com/dvoelkl/HIDE-deconv).

### 3.2 Benchmark

We evaluated HIDE-Deconv using benchmark datasets generated from the independent LUAD test partition described in Section S1.2.1. Because the true cellular compositions of the resulting pseudobulk samples are known, they provide a controlled setting for evaluating deconvolution performance across multiple levels of cellular resolution. The mean prediction accuracy and the corresponding standard deviation for the ten independent benchmark runs are provided in Supplementary Tables S10-S12. To facilitate reproducibility, complete cell-type-specific benchmark results, benchmark workflows and analysis scripts are provided through Zenodo (DOI: 10.5281/zenodo.24117378).

#### 3.2.1 HIDE-Deconv achieves the highest overall benchmark performance

Predictive performance was evaluated by comparing the true cellular compositions of the pseudobulk samples (*C*) with the cellular compositions estimated by each deconvolution method (*C*^^^). Methods capable of operating across multiple hierarchy levels were evaluated at all supported levels of cellular resolution. For methods without native hierarchical support, major-cell-type abundances were additionally obtained through aggregation of sub-minor cell-type predictions according to the cell-type hierarchy.

Across hierarchy levels, HIDE-Deconv achieved consistently strong performance, yielding high Pearson and Spearman correlations together with low NMAE values (Figure 2A-C, Supplementary Tables S10-S12, including standard deviations). At major-cell-type resolution, HIDE-Deconv, HIDE and DTD consistently ranked among the best-performing methods. HIDE-Deconv achieved the strongest performance in most benchmark categories and was particularly competitive at sub-minor-cell-type resolution. Rectangle and BayesPrism represented the strongest competing approaches. Rectangle achieved favorable NMAE values at minor- and sub-minor-cell-type resolution, whereas BayesPrism ranked among the strongest methods at finer levels of cellular resolution. In contrast, PREDE, MuSiC and EPIC generally showed lower performance across benchmark metrics.

**Fig 2.**
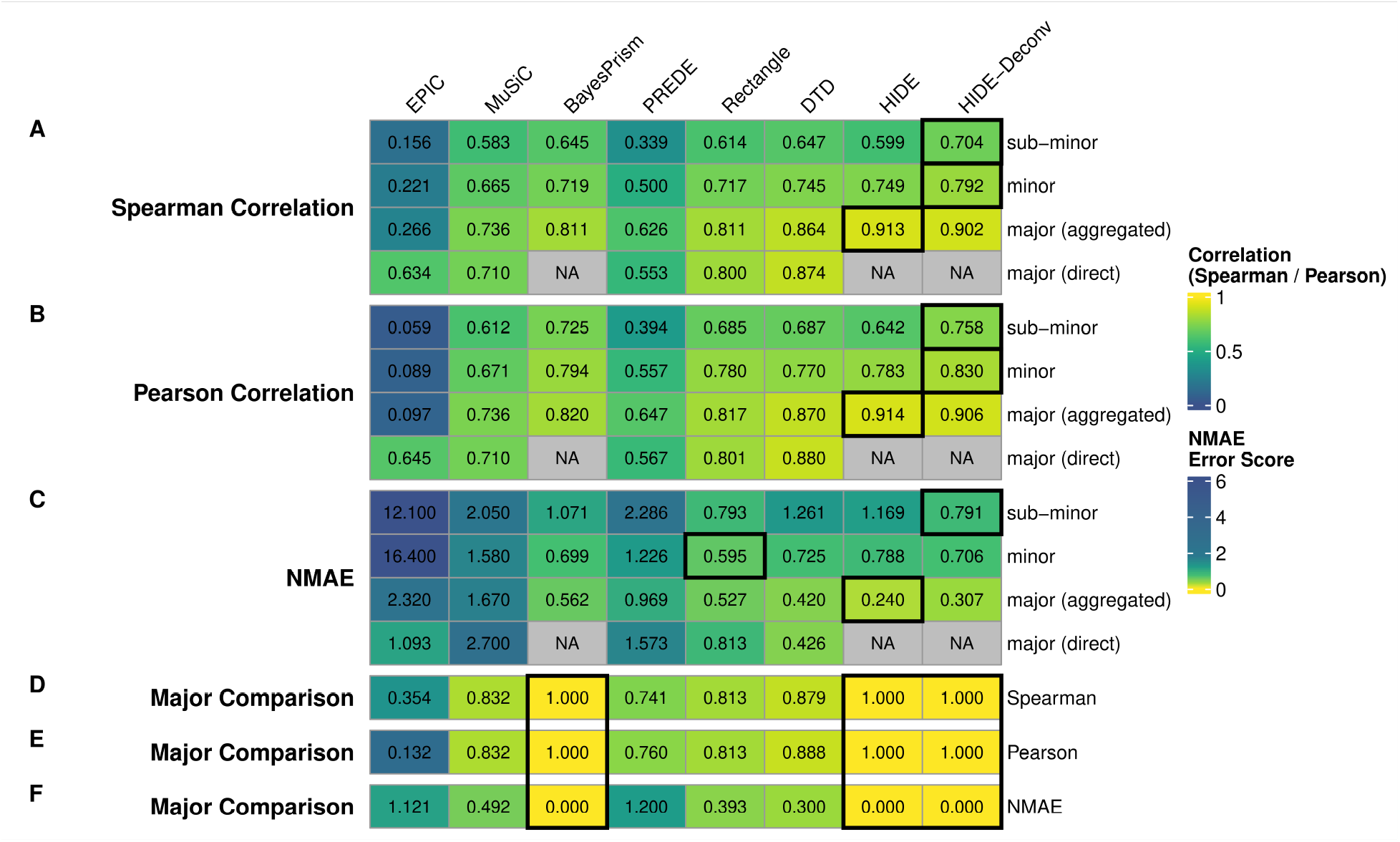
Benchmark performance and hierarchical consistency of HIDE-Deconv and competing deconvolution methods. (A-C) Benchmark performance measured by Spearman correlation (A), Pearson correlation (B), and normalized mean absolute error (NMAE) (C). Metrics are reported for sub-minor cell types, as well as for minor- and major-cell-type abundances obtained by hierarchical aggregation of sub-minor cell-type predictions. For methods without native hierarchical support, major-cell-type abundances were additionally estimated directly at the major-cell-type level. The values represent averages across all cell types within a given hierarchy level. The best-performing method for each benchmark metric and hierarchy level is highlighted by a black border. (D-F) Hierarchical consistency measured by Spearman correlation (D), Pearson correlation (E), and NMAE (F). Major-cell-type abundances reconstructed by aggregating sub-minor cell-type predictions were compared with major-cell-type abundances estimated directly at the major-cell-type level. These metrics quantify the extent to which a method produces consistent cellular compositions across hierarchy levels. Higher correlation values and lower NMAE values indicate greater agreement between both estimation strategies and therefore greater hierarchical consistency. The benchmark samples consisted of pseudobulk RNA-seq profiles generated from the LUAD single-cell dataset (see Supplementary Method Section S1.4.2). NA indicates that no corresponding value was available because the respective hierarchical method inferred major-cell-type abundances through aggregation of finer-resolution predictions and therefore did not generate an independent direct major-cell-type estimate. Standard deviations are provided in Supplementary Tables S10-S12.

Performance decreased from major- to minor- and sub-minor-cell-type resolution for all methods, reflecting the increasing difficulty of resolving closely related cellular populations. Nevertheless, HIDE-Deconv maintained strong predictive performance across all hierarchy levels. The largest advantages relative to competing methods were observed at minor- and sub-minor-cell-type resolution, where accurate discrimination between closely related cellular populations becomes most challenging.

For non-hierarchical methods, major-cell-type abundances were evaluated both by direct prediction at the major-cell-type level (*major (direct)*) and by aggregation of sub-minor cell-type predictions. Interestingly, major-cell-type abundances reconstructed from sub-minor predictions generally achieved higher predictive performance than abundances estimated directly at the major-cell-type level, with EPIC representing the main exception.

#### 3.2.2 HIDE-Deconv provides consistent estimates across hierarchy levels

To evaluate hierarchical consistency, we compared major-cell-type abundances obtained by aggregating sub-minor predictions with abundances estimated directly at the major-cell-type level (Figure 2D-F). Methods with native hierarchical inference procedures (HIDE, BayesPrism and HIDE-Deconv) are hierarchically consistent by construction.

Among the methods without explicit hierarchical modeling, DTD achieved the highest consistency (Spearman = 0.879, Pearson = 0.888, NMAE = 0.300), followed by Rectangle. PREDE, MuSiC and EPIC showed substantially larger discrepancies between aggregated and directly estimated major-cell-type abundances, with EPIC showing the weakest agreement overall. While MuSiC achieved moderate Pearson and Spearman correlations, its high NMAE indicates substantial deviations in absolute abundance estimates.

#### 3.2.3 HIDE-Deconv combines the strengths of DTD and HIDE

To assess the contributions of hierarchical modeling and gene-weight optimization, we compared HIDE-Deconv with its predecessor methods DTD and HIDE. DTD does not explicitly incorporate hierarchical relationships between cell types, whereas HIDE performs hierarchy-aware deconvolution through a top-down inference strategy. HIDE-Deconv combines both concepts within a joint optimization framework.

HIDE-Deconv achieved better performance across hierarchy levels compared with DTD and HIDE despite for HIDE outperforming slightly at major level while HIDE-deconv additionally maintaining hierarchical consistency.

In contrast, HIDE with its top-down strategy generally achieved strong performance at major-cell-type resolution but showed reduced performance at finer levels of cellular resolution.

Overall, HIDE-Deconv combines the hierarchical consistency introduced by HIDE with increased predictive performance comparable to DTD, while providing a more balanced performance profile across multiple levels of cellular resolution.

### 3.3 Hierarchy-informed deconvolution links global tissue remodeling to fine cell-type changes during lung cancer progression

To assess the biological utility of hierarchical deconvolution within a complete HIDE-Deconv workflow, we analyzed bulk RNA-seq data from 110 lung adenocarcinoma tumors collected prior to PD-L1 inhibitor treatment ((23)). For deconvolution, HIDE-Deconv was trained on the complete LUAD single-cell reference dataset described in Methods Section 2.2 using a three-level hierarchy comprising 10 major, 17 minor and 32 sub-minor cell-type populations (Supplementary Table S5). We subsequently compared cellular compositions between early-stage (I+II) and late-stage (III+IV) tumors (patient demographics in Supplementary Table S13).

At the major-cell-type level, late-stage tumors exhibited a marked shift from epithelial-dominated toward more immune-enriched tissue compositions (Figure 3). Epithelial-cell proportions decreased substantially (0.60 vs. 0.30, *p*_adj_ *<* 0.001), whereas several immune-cell populations increased, including T-cells (0.11 vs. 0.23, *p*_adj_ *<* 0.001), NK-cells (0.02 vs. 0.05, *p*_adj_ = 0.01) and B-cells (0.02 vs. 0.07, *p*_adj_ = 0.05). A corresponding heatmap visualization across major-, minor- and sub-minor-cell-type populations is provided in Supplementary Figure S1.

**Fig 3.**
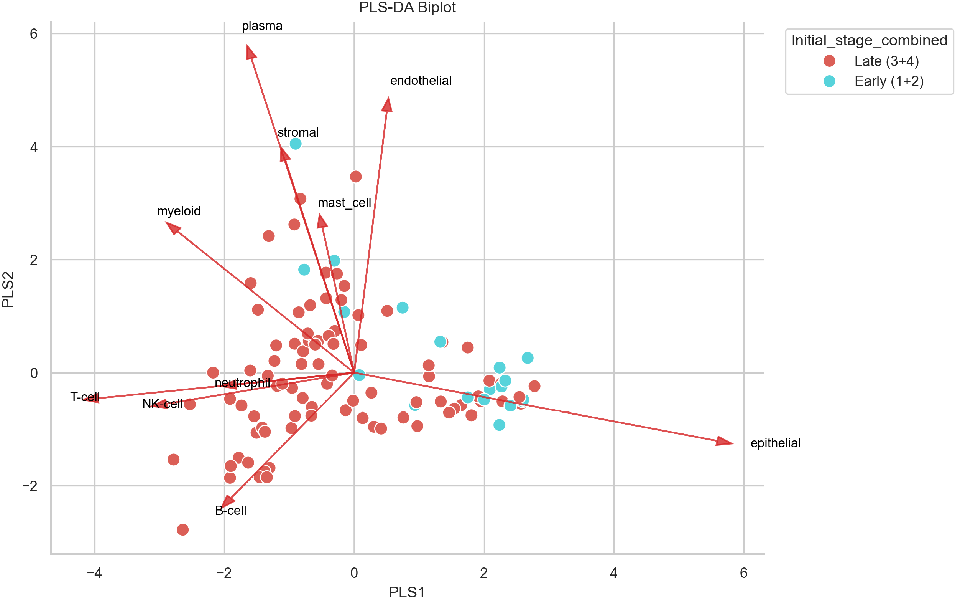
Global tissue remodeling during LUAD progression at major-cell-type level. Partial least-squares discriminant analysis (PLS-DA, see Supplementary Methods Section S1.5) of major-cell-type compositions estimated by HIDE-Deconv in early-stage (I+II) and late-stage (III+IV) LUAD tumors. The arrows indicate the contributions of individual major cell types to the separation between the two groups. Separation along the first latent component reflects differences in overall tissue composition, characterized by higher epithelial-cell proportions in early-stage tumors and increased immune-cell proportions in late-stage tumors.

While the major-cell-type analysis revealed a reduction in epithelial-cell abundance, hierarchical deconvolution provided a more detailed view of the underlying epithelial subpopulations. At the minor-cell-type level, the alveolar compartment decreased substantially (0.39 vs. 0.14, *p*_adj_ = 0.002), while club-cell abundance was also reduced (0.05 vs. 0.02, *p*_adj_ = 0.02).

Although the alveolar compartment decreased overall, sub-minor-cell-type analysis revealed opposing trends among its constituent populations that were not apparent at broader levels of cellular resolution. Pulmonary alveolar type 1 cells increased (0.001 vs. 0.006, *p*_adj_ = 0.02), whereas pulmonary alveolar type 2 cells decreased (0.39 vs. 0.13, *p*_adj_ = 0.003) with advancing disease stage.

The increase in T-cell abundance could likewise be traced to specific T-cell populations at finer levels of resolution, including CD8+ alpha-beta T-cells (0.05 vs. 0.13, *p*_adj_ = 0.04) and regulatory T-cells (0.02 vs. 0.03, *p*_adj_ = 0.04). In contrast, no significant differences were observed for the major myeloid compartment. However, hierarchical analysis revealed increased monocyte abundance in late-stage tumors (0.03 vs. 0.07, *p*_adj_ = 0.02), predominantly driven by classical monocytes (0.03 vs. 0.06, *p*_adj_ = 0.02).

Additional significantly altered immune-cell populations are reported in Supplementary Tables S14-S16, which provide the complete results of all Mann-Whitney U tests across all hierarchy levels.

### 3.4 Hierarchical gene weighting reveals progressively refined cell-type signatures

To investigate how hierarchical gene weighting resolves lineage- and cell-type-specific molecular signatures, we examined genes prioritized by HIDE-Deconv during training on the LUAD single-cell reference dataset. Genes were ranked according to a score combining learned gene weights and expression variability (Supplementary Methods Section S1.5.2), enabling comparison of informative features across hierarchy levels.

At the major-cell-type level, the highest-ranking genes reflected established lineage-associated signatures across epithelial, immune, stromal, endothelial and B-cell populations (Figure 4 a)). Stromal-associated genes including TIMP1, COL6A2, LUM, BGN and COL1A1 formed a characteristic lineage-specific signature.

**Fig 4.**
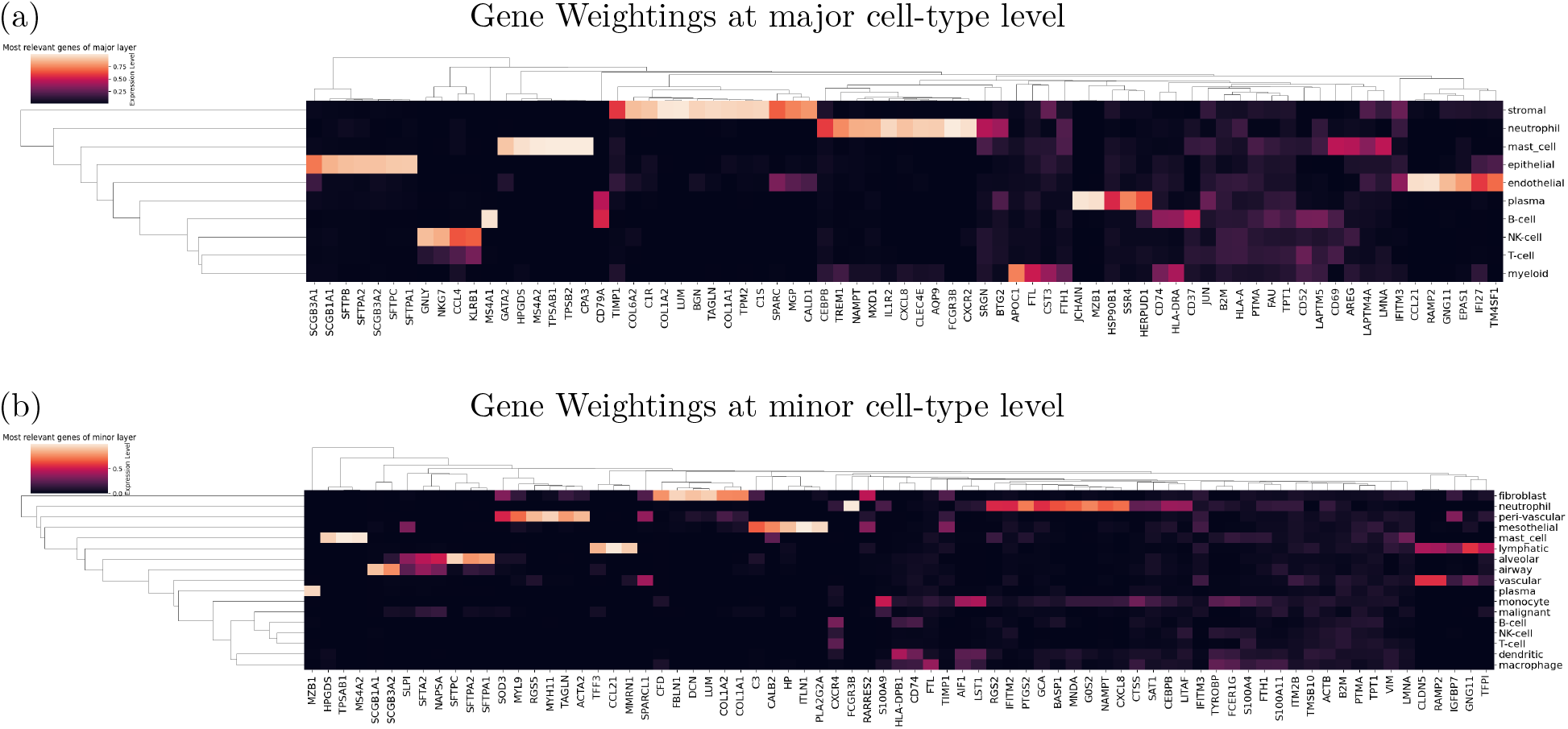
Hierarchy-specific gene signatures learned by HIDE-Deconv. Major-cell-type (a) and minor-cell-type (b) marker heatmaps derived from learned gene weights. Genes were ranked according to the score *s_i_* = *g_i_* · Var(*X_i_*) (Supplementary Methods Section S1.5.2) and the 75 highest-ranked genes are shown. Gene expression values were scaled between 0 and 1 for visualization. While broad lineage-associated signatures distinguish major cellular compartments, finer hierarchy levels progressively resolve more specific cell-type signatures within these populations.

To assess how gene specificity evolves across hierarchy levels, we focused on the fibroblast lineage as a representative stromal compartment. While fibroblast-associated genes were already apparent at the major-cell-type level, additional markers emerged at the minor-cell-type level, resulting in progressively refined cell-type signatures (Figure 4 b)).

At sub-minor resolution, these signatures were further resolved into subtype-specific patterns. For example, COL1A2 and COL3A1 showed selective enrichment in bronchial fibroblasts, distinguishing them from other fibroblast populations (Supplementary Figure S2).

### 3.5 Tumor microenvironment states are associated with response to PD-L1 blockade

We next investigated whether hierarchical deconvolution could identify distinct tumor microenvironment (TME) states that were not explained solely by clinical stage. Samples were clustered based on sub-minor cell-type abundances, resulting in three well-separated TME states (Figure 5). Alternative clustering solutions are shown in Supplementary Figures S3-S5.

**Fig 5.**
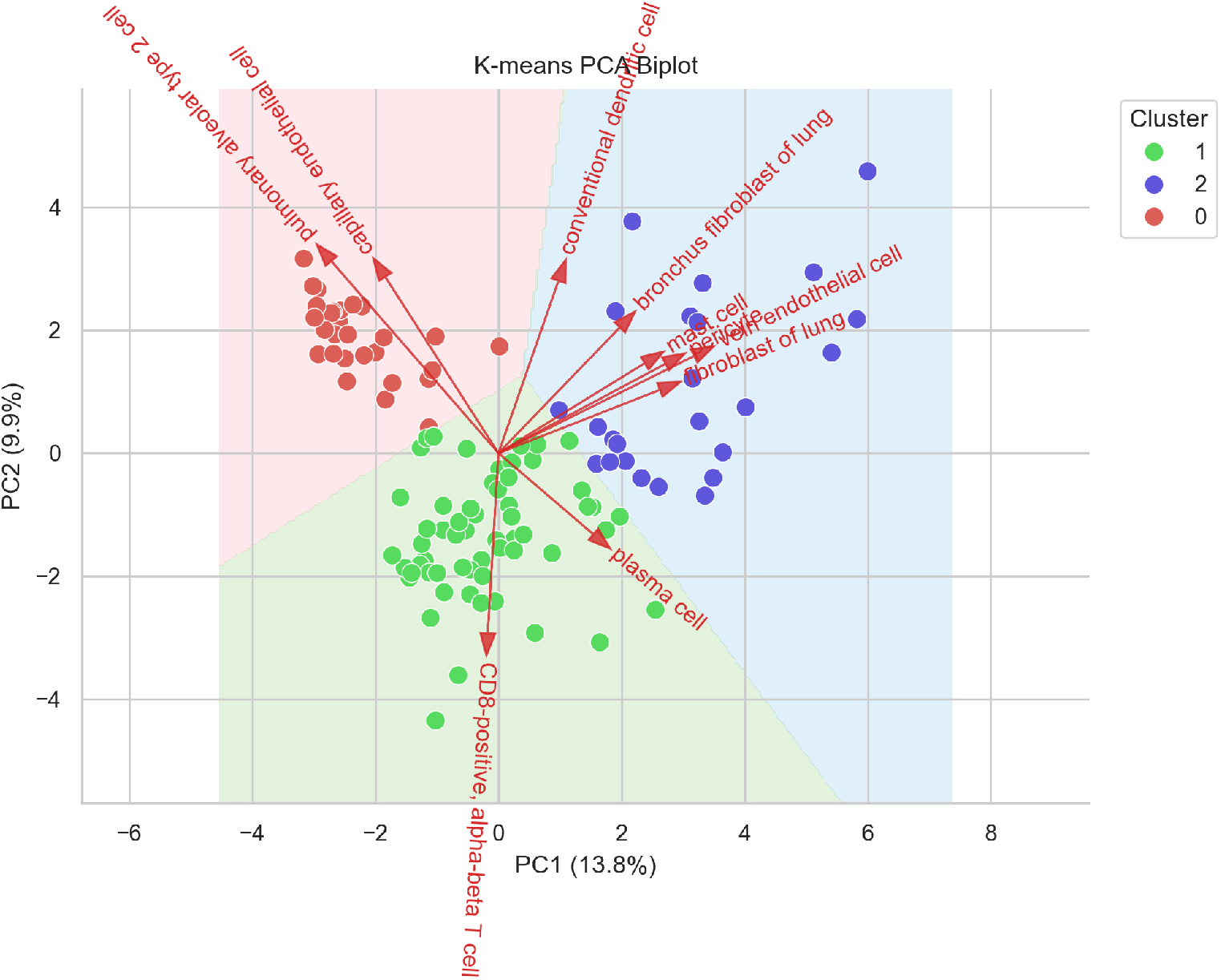
Three tumor microenvironment states identified from deconvolved cellular compositions. PCA biplot of LUAD samples based on sub-minor cell-type abundances. Samples are colored according to the three identified clusters. Arrows indicate cell populations contributing to cluster separation.

Principal component analysis demonstrated clear separation between the three clusters, indicating substantial differences in cellular composition. To characterize the resulting TME states, cell-type abundances were compared across all hierarchy levels using Kruskal-Wallis tests followed by post-hoc Dunn tests (Supplementary Tables S15-S17).

#### 3.5.1 Hierarchical deconvolution identifies three distinct TME states

Comparison of major cell-type abundances revealed one epithelial-dominated cluster and two immune-enriched clusters. At the major-cell-type level, clusters 1 and 2 appeared largely similar, as both exhibited elevated immune-cell abundances. However, hierarchical analysis at higher cellular resolution revealed substantial differences in the composition of the underlying immune microenvironment. Cluster 0 was characterized by increased epithelial-cell abundance, whereas clusters 1 and 2 both showed increased lymphoid and immune-cell infiltration. Cluster 2 additionally exhibited elevated stromal and endothelial-cell proportions, suggesting a distinct stromal-immune microenvironment state (see Figure 5).

Hierarchical deconvolution further resolved the cellular populations underlying these major-cell-type patterns (Supplementary Table 15). The epithelial-dominated cluster 0 was characterized by increased alveolar type 2 cells (*ρ*_0_ = 0.57, *ρ*_1_ = 0.05, *ρ*_2_ = 0.02, *p*_adj_ *<* 0.001) and increased malignant-cell proportions. In contrast, cluster 1 showed a lymphocyte-rich composition with elevated CD8+ T-cells (*ρ*_0_ = 0.05, *ρ*_1_ = 0.17, *ρ*_2_ = 0.07, *p*_adj_ *<* 0.01) together with increased B-cell, NK-cell and regulatory T-cell abundances (Supplementary Table 17).

Although cluster 2 also exhibited elevated immune-cell abundances at the major-cell-type level, analysis at finer levels of resolution revealed a distinct composition characterized by increased myeloid, stromal and endothelial-cell populations. Increased classical monocyte abundance (*ρ*_0_ = 0.009, *ρ*_2_ = 0.049, *p*_adj_ *<* 0.001) and fibroblast abundance (*ρ*_0_ = 0.002, *ρ*_2_ = 0.018, *p*_adj_ *<* 0.001) distinguished cluster 2 from both the epithelial-dominated cluster 0 and the lymphocyte-rich cluster 1.

Additional significantly altered cell populations are reported in Supplementary Tables S15-S17.

Comprehensive heatmap visualizations of cell-type abundances across the three TME states are provided in Supplementary Figures S6-S8.

#### 3.5.2 Tumor microenvironment states are associated with distinct transcriptional programs

To determine whether the TME states identified by HIDE-Deconv were associated with distinct transcriptional programs, we compared gene expression across the three clusters using differential expression analysis. Genes upregulated in each pairwise cluster comparison were subsequently analyzed for enrichment of Cancer Hallmark gene sets using the platform described by Menyhart et al. (26) to characterize the biological programs associated with the respective TME states (Supplementary Methods Section S1.5.3).

No Cancer Hallmark reached statistical significance when comparing clusters 0 and 1, despite substantial differences in cellular composition identified by hierarchical deconvolution. In contrast, comparison of the epithelial-dominated cluster 0 with the stromal-immune cluster 2 revealed significant enrichment of pathways associated with tissue invasion and metastasis in cluster 2 (Supplementary Figure S9).

Although clusters 1 and 2 both displayed elevated immune-cell abundances at the major-cell-type level, hierarchical deconvolution revealed two distinct immune microenvironment states. Relative to cluster 1, cluster 2 was enriched for transcriptional programs associated with tissue invasion and metastasis. In contrast, cluster 1 showed enrichment of Hallmarks related to evading immune destruction and tumor-promoting inflammation (Figure 6).

**Fig 6.**
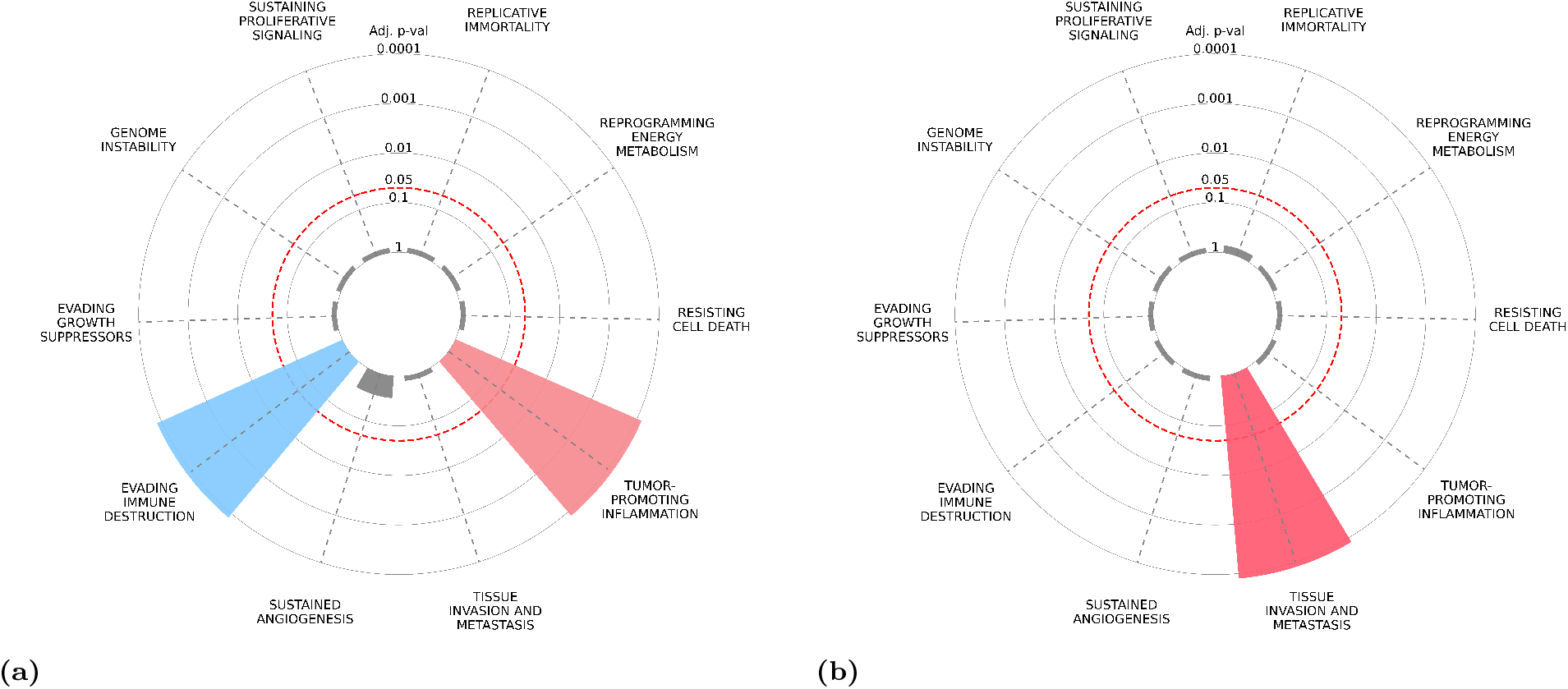
Distinct transcriptional programs characterize two immune-enriched tumor microenvironment states. Cancer Hallmark enrichment analysis comparing the lymphocyte-rich TME state (cluster 1) and the stromal-immune TME state (cluster 2). The left panel shows Hallmarks enriched in cluster 1, whereas the right panel shows Hallmarks enriched in cluster 2. Hallmarks outside the red circle reached statistical significance.

#### 3.5.3 Distinct tumor microenvironment states are associated with PD-L1 response and survival

Given that hierarchical deconvolution distinguished two immune-enriched TME states with distinct transcriptional programs, and considering the established role of the tumor microenvironment in determining response to PD-L1 blockade (27), we next investigated whether the identified TME states were associated with treatment response. Patients assigned to cluster 1 exhibited a significantly higher probability of responding to treatment than patients assigned to cluster 2 (relative risk 3.00, 95% CI 1.23 − 7.34, *p* = 0.016). In contrast, patients assigned to the epithelial-dominated cluster 0 showed similar numbers of responders and non-responders (10 vs. 10 patients; Supplementary Table S20).

We next examined whether these response differences were reflected in clinical outcome. Consistent with the response analysis, patients assigned to cluster 1 showed improved 6-month overall survival compared with cluster 2 (Figure 7 a)). Similarly, all responders remained alive during the first six months of follow-up, whereas survival among non-responders fell below 60% during the same period (Figure 7 b)).

**Fig 7.**
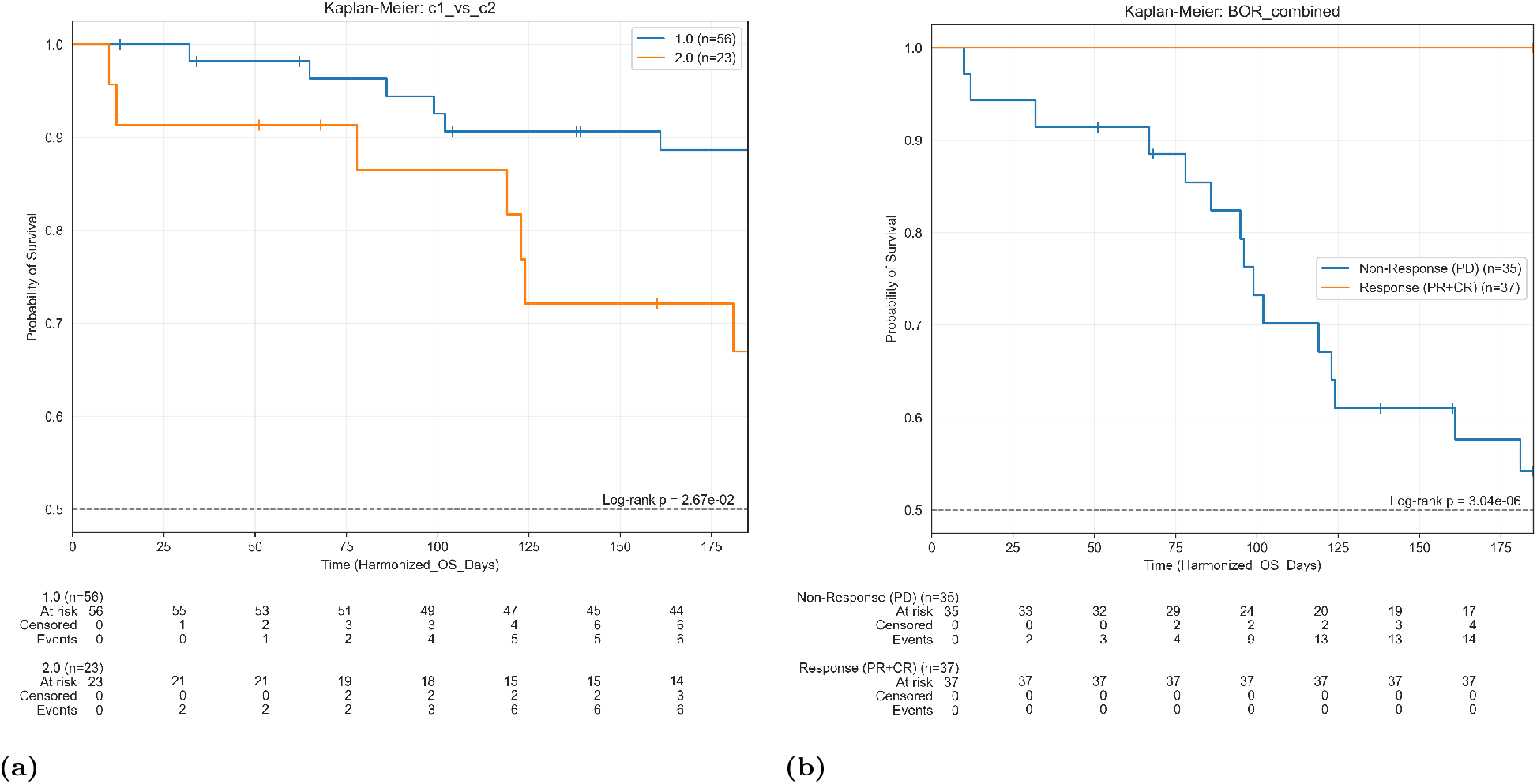
Kaplan-Meier analysis of 6-month overall survival. (A) Survival stratified by tumor microenvironment (TME) state. Patients assigned to the lymphocyte-rich cluster 1 showed improved survival compared with patients assigned to the stromal- and myeloid-enriched cluster 2. (B) Survival stratified by treatment response. Responders exhibited improved survival compared with non-responders, and all responders remained alive during the first six months of follow-up. Therapy-response data were not available for all patients.

### 3.6 Disease-associated immune remodeling becomes apparent at different levels of cellular resolution

To investigate whether hierarchical cellular remodeling extends beyond the tumor microenvironment, we applied HIDE-Deconv to PBMC-derived bulk RNA-seq datasets from patients with sepsis, COVID-19 and systemic lupus erythematosus (SLE). For all analyses, deconvolution was performed using a common PBMC reference dataset and a two-level hierarchy comprising 7 major and 25 minor cell-type populations. Detailed dataset descriptions and preprocessing procedures are provided in the Supplementary Methods S1.2.2 and Supplementary Tables S6-S8.

To compare how disease-associated immune remodeling manifests across cellular hierarchies, we performed PLS-DA on deconvolved major- and minor-cell-type abundances (Figure 8). The degree of separation between patients and healthy controls differed substantially between diseases and hierarchy levels.

**Fig 8.**
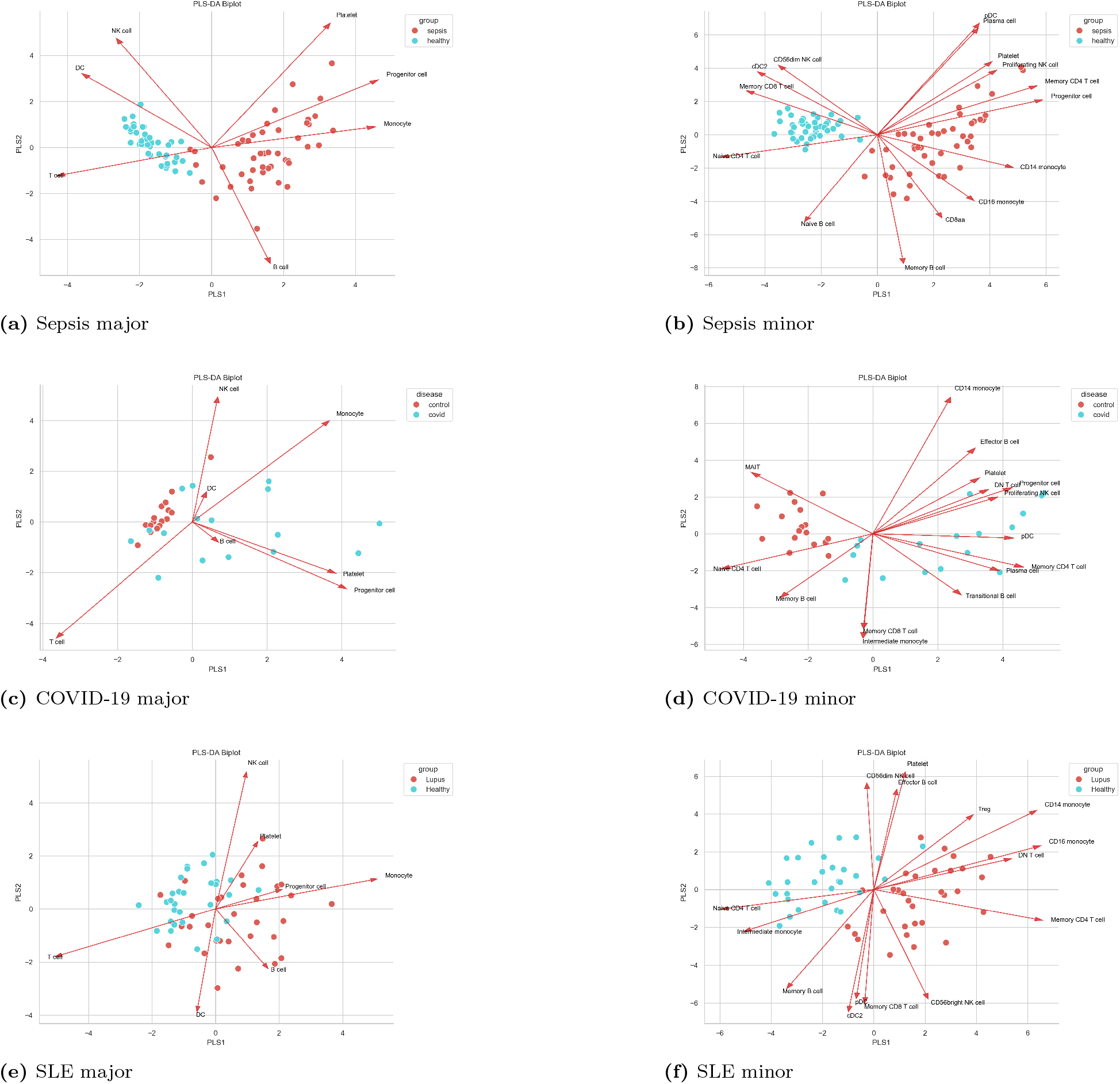
PLS-DA of major- and minor-cell-type compositions across PBMC disease cohorts. PLS-DA was performed on HIDE-Deconv-derived cellular compositions from sepsis (A-B), COVID-19 (C-D) and SLE (E-F) cohorts. Left panels show major-cell-type compositions and right panels show minor-cell-type compositions. Arrows indicate cell populations contributing to group separation. While sepsis samples were already well separated at the major-cell-type level, separation between patients and controls became more pronounced at minor-cell-type resolution in COVID-19 and SLE, illustrating that disease-associated immune remodeling can become apparent at different levels of cellular resolution.

In sepsis, patient and control samples were already almost completely separated at the major-cell-type level, indicating that disease-associated remodeling is dominated by broad lineage-level changes. Major shifts included increased monocyte abundance (0.31 vs 0.09, *p*_adj_ = 2.42 × 10*^−^*^11^) together with reduced T-cell (0.52 vs 0.71, *p*_adj_ = 7.35 × 10*^−^*^11^), NK-cell (0.05 vs 0.10, *p*_adj_ = 1.49 × 10*^−^*^5^) and dendritic-cell proportions (0.008 vs 0.030, *p*_adj_ = 3.02 × 10*^−^*^11^). Minor-cell-type analysis provided additional resolution but largely refined patterns that were already apparent at the major-cell-type level (Supplementary Tables S21-S22, Supplementary Figure S10).

In contrast, COVID-19 showed comparatively limited changes at the level of major immune-cell lineages. Significant changes at major-cell-type resolution were largely restricted to progenitor-cell (0.0021 vs 0.0000, *p*_adj_ = 3.37 × 10*^−^*^5^) and platelet abundances (0.0020 vs 0.0000, *p*_adj_ = 0.0011). However, analysis at minor-cell-type resolution revealed marked differences involving plasma cells (0.027 vs 0.0000, *p*_adj_ = 1.07 × 10*^−^*^5^), naive and memory CD4+ T-cells, proliferating NK-cells and plasmacytoid dendritic cells (Supplementary Tables S23-S24, Figure 8). In contrast to the major-cell-type analysis, substantial differences between patients and controls became apparent across multiple immune-cell populations at minor-cell-type resolution.

Systemic lupus erythematosus (SLE) displayed an intermediate pattern. Major-cell-type analysis identified significant alterations in T-cell (0.58 vs 0.67, *p*_adj_ = 0.0108) and monocyte abundances (0.23 vs 0.15, *p*_adj_ = 0.0011), while minor-cell-type analysis revealed additional changes involving memory CD4+ T-cells, double-negative T-cells, regulatory T-cells and multiple monocyte populations (Supplementary Tables S25-S26, Supplementary Figure S11).

These patterns were also evident in the PLS-DA analyses shown in Figure 8. In sepsis, patient and control samples were already well separated using major-cell-type compositions, whereas COVID-19 samples showed separation only clearly at minor-cell-type resolution. Samples from patients with systemic lupus erythematosus were separated at both hierarchy levels, with more pronounced separation observed at the minor-cell-type level.

## 4 Discussion

The increasing availability of large single-cell reference atlases has substantially expanded the scope of bulk RNA-seq deconvolution (8; 28; 29). In addition, many current reference atlases provide cell-type annotations at multiple levels of cellular resolution, ranging from broad cellular lineages to highly specialized cellular subpopulations. However, most existing approaches still estimate cellular compositions at a single annotation level or treat different resolutions independently. Our results demonstrate that explicitly incorporating hierarchical cell-type relationships can improve both the consistency of deconvolution estimates and the biological interpretation of disease-associated cellular remodeling.

The benchmark results demonstrate that cellular compositions estimated at different hierarchy levels are not necessarily mutually consistent. Major-cell-type abundances obtained by direct deconvolution frequently differed from abundances obtained by aggregating fine-grained predictions, despite both estimates describing the same biological quantity. Systematic evaluation of hierarchical consistency revealed these discrepancies across multiple deconvolution approaches. By explicitly incorporating hierarchical cell-type relationships into a joint optimization framework, HIDE-Deconv improves hierarchical consistency while maintaining predictive performance across multiple levels of cellular resolution.

By preserving consistency between broad cellular lineages and their constituent subpopulations, HIDE-Deconv facilitates high-resolution cell-type deconvolution and enables cellular remodeling to be investigated across multiple levels of biological organization within a unified analytical framework.

The benchmark also provides insight into the performance differences between HIDE-Deconv and its predecessor HIDE. The lower performance of HIDE in the present benchmark compared with the original publication can largely be explained by differences in the benchmarking strategy. Unlike the original study, which used cell-level partitioning, the present benchmark employed a study-level train-test split. This strategy provides a more stringent assessment of generalization by evaluating performance across independent studies rather than across cells originating from the same dataset. Despite this substantially more challenging benchmark setting, HIDE-Deconv maintained predictive performance while ensuring consistency across hierarchy levels.

In addition to cellular abundance estimation, HIDE-Deconv learns hierarchy-specific gene signatures through the joint optimization of hierarchy-specific gene weights. The resulting markers recapitulated established lineage-associated gene-expression patterns while progressively refining cell-type-specific signatures at finer levels of cellular resolution, suggesting that the learned weights capture biologically meaningful features of the underlying cellular hierarchy.

This principle is illustrated by clusters 1 and 2 of the LUAD tumor microenvironment analysis, where two TME states exhibited similarly high overall immune-cell abundances at the major-cell-type level yet differed substantially in their cellular composition at finer levels of resolution. Although both states were characterized by elevated immune-cell abundances, they exhibited distinct transcriptional programs. One showed enrichment of hallmarks related to tissue invasion and metastasis, whereas the other was enriched for hallmarks associated with evading immune destruction and tumor-promoting inflammation. Moreover, the two immune-enriched states differed not only in cellular composition but also in treatment response and associated clinical outcome, highlighting biological heterogeneity that remained unresolved at the major-cell-type level.

While the LUAD analyses demonstrate the utility of hierarchical deconvolution in solid tumors, the PBMC analyses indicate that the same concept generalizes to systemic immune responses. Together, these analyses suggest that hierarchical cell-type remodeling is not restricted to the tumor microenvironment and can also be observed across diverse inflammatory and autoimmune diseases.

Different diseases revealed disease-associated remodeling at different levels of the hierarchy. While major-cell-type abundances captured most of the sepsis signal, COVID-19-associated changes became apparent primarily at minor-cell-type resolution, whereas SLE showed alterations at both levels. Examining multiple hierarchy levels therefore provides information that would be missed when analyses are restricted to a single cellular resolution.

Hierarchical approaches provide a means of studying cellular remodeling across multiple cellular scales without restricting analyses to a predefined level of resolution. Whereas some diseases are characterized primarily by broad lineage-level shifts, others exhibit remodeling that only becomes apparent within specific cellular subpopulations. By linking major cellular compartments to their constituent subpopulations, hierarchical deconvolution facilitates the interpretation of compositional changes and helps distinguish global tissue-level shifts from redistribution within individual cellular lineages. More generally, changes observed at broad cellular resolutions can be traced back to the populations that contribute to them, providing a common framework for comparing remodeling patterns across different diseases and tissues.

This principle was consistently observed across the PBMC analyses. In COVID-19, most disease-associated alterations became apparent only at minor-cell-type resolution despite comparatively limited changes at the level of major immune-cell lineages. In contrast, sepsis was characterized by pronounced lineage-level remodeling involving monocyte and T-cell populations, whereas SLE combined broad lineage-level shifts with extensive subtype-specific changes. These results indicate that the cellular resolution required to capture disease-associated remodeling differs across diseases. Examining multiple hierarchy levels therefore provides a more complete view of disease-associated changes than analyses restricted to a single cellular scale.

Beyond improving hierarchical deconvolution itself, HIDE-Deconv addresses practical barriers that frequently limit the adoption of deconvolution methods in biomedical research. HIDE-Deconv integrates reference preparation, model training, hierarchical deconvolution and downstream statistical analyses within a unified and reproducible workflow. Using annotated single-cell reference data, bulk RNA-seq expression profiles and optional clinical metadata, the framework supports end-to-end hierarchical deconvolution analyses through a single command-line interface. By reducing workflow fragmentation and technical overhead, HIDE-Deconv facilitates the application of high-resolution cell-type deconvolution in translational and biomedical research.

Like other reference-based deconvolution methods, HIDE-Deconv remains dependent on the quality and availability of suitable single-cell reference data (8; 28). Cellular populations that are absent from the reference may lead to inaccurate abundance estimates for related cell types. In addition, disease-associated transcriptional changes may occasionally resemble compositional shifts when cellular populations deviate substantially from their reference profiles. Although HIDE-Deconv applies gene-median normalization to reduce differences between single-cell and bulk RNA-seq platforms, accurate estimation of cross-platform correction factors depends on the availability of bulk RNA-seq samples representative of the analyzed dataset.

Several recent methodological developments suggest possible directions for addressing these limitations.

Approaches based on matrix factorization have demonstrated the ability to infer previously unmodeled cellular populations directly from bulk transcriptomic data (30; 31). Other methods have addressed hidden cellular contributions and cell-type-specific gene regulation (32), while approaches such as Harp were specifically developed to improve robustness across sequencing platforms (33). Future extensions of HIDE-Deconv may benefit from integrating such concepts into hierarchical deconvolution frameworks.

## 5 Conclusion

HIDE-Deconv extends hierarchical deconvolution by jointly optimizing predictions across multiple levels of cellular resolution while maintaining consistency between hierarchy levels. In benchmark analyses, HIDE-Deconv achieved strong predictive performance across hierarchy levels and improved performance relative to the sequential hierarchical strategy employed by HIDE.

Application to lung adenocarcinoma and PBMC datasets showed that disease-associated cellular remodeling can become apparent at different levels of a cellular hierarchy. While some biological signals were already captured at broad lineage-level resolution, others emerged only when specific cellular subpopulations were considered. Examining multiple hierarchy levels therefore provides a more complete view of disease-associated cellular changes than analyses restricted to a single level of cellular resolution.

In addition to the hierarchical model itself, HIDE-Deconv provides an open-source framework that integrates reference construction, model training, deconvolution and downstream analyses within a unified workflow. We hope that this combination of hierarchical modeling and accessible software will facilitate the broader use of hierarchical deconvolution approaches in biomedical research.

## Supporting information

**S1 Appendix.** Supplementary Methods, Supplementary Figures S1–S12 and Supplementary Tables S1–S24.

**Software availability.** HIDE-Deconv is available through PyPI:

https://pypi.org/project/hide-deconv/ and GitHub: https://github.com/dvoelkl/HIDE-deconv.

**Benchmark scripts and reproducibility material.** Zenodo DOI: 10.5281/zenodo.24117378

## Acknowledgments

We thank all contributors of the publicly available datasets used in this study.

