## Supplement_1 for "Characterizing multiscale cellular remodeling through hierarchical deconvolution with HIDE-Deconv"

### Supplementary Material

**2 Cell Type Distribution and hierarchy of PBMC Dataset** **10**

**3 Bulk Datasets** **13**

**4 Results Benchmark** **15**

### 1 Supplementary Methods

#### 1.1 Digital Tissue Deconvolution

Feature selection strongly influences the performance of transcriptomic deconvolution methods. In our previous work DTD(1), we showed that continuously optimized gene weights learned from annotated single-cell training data can substantially improve deconvolution accuracy. HIDE-Deconv builds upon this weighted deconvolution framework and extends it to hierarchical cellular annotations. We first introduce the underlying weighted least-squares formulation before describing its hierarchical extension.

Let $X \in \mathbb{R}^{p \times q}$ denote the matrix of $q$ cellular reference profiles, with each column representing one cell-type-specific reference profile, and let $Y \in \mathbb{R}^{p \times n}$ denote the matrix of $n$ bulk gene expression profiles. The rows of both matrices correspond to the same set of $p$ genes.

In DTD, bulk expression profiles are represented as linear combinations of the reference profiles weighted by the cellular composition matrix $C \in \mathbb{R}^{q \times n}$,

$$Y = XC. \tag{1}$$

Each column of $C$ corresponds to the cellular composition of a single sample. If all cell types present in the bulk sample are represented in $X$, the corresponding proportions sum up to 1.
A naive estimate of $C$ can be obtained by solving the least-squares problem

$$\hat{C} = \arg\min_{C} ||Y - XC||_F^2, \tag{2}$$

where $\|\cdot\|_F$ denotes the Frobenius norm, the standard matrix norm obtained by summing squared residuals across all genes and samples. Given $X$ and $Y$, $\hat{C}$ contains the estimated cellular compositions.

The least-squares formulation can be extended by introducing gene-specific weights $g = (g_1, g_2, ..., g_p)^T$ by

$$\hat{C}(g) = \arg\min_{C} ||\text{diag}(g)(Y - XC)||_F^2 \tag{3}$$

subject to the element-wise non-negativity constraint

$$\hat{C}(g) \geq 0 \tag{4}$$

The gene weights are learned by minimizing an outer objective function defined on bulk samples with known cellular compositions:

$$L(g) = -\sum_{j=1}^{q} \text{corr}(C_{j,\cdot}, \hat{C}_{j,\cdot}(g)) \text{ subject to } ||g||_2 = 1, \tag{5}$$

with corr$(\cdot, \cdot)$ denoting the Pearson correlation. Pearson correlation was selected because the objective function aims to maximize agreement between estimated and true cellular proportions on a continuous scale rather than only their relative ranking. The objective function is minimized when the learned gene weights maximize the agreement between estimated and true cellular compositions, thereby improving the deconvolution performance of the weighted least-squares model.

#### 1.2 Reference datasets

##### 1.2.1 LUAD single-cell reference dataset

As a single-cell reference for benchmark generation and lung adenocarcinoma analyses, we used the extended single-cell lung cancer atlas (LuCA) published by Salcher et al. (2). This LUAD reference dataset was obtained from the CZ CELLxGENE Discover portal (3). The original atlas integrates multiple previously published lung cancer single-cell RNA-seq studies and contains both malignant and non-malignant lung samples. For this work, we restricted the dataset to primary LUAD tumor samples originating from lung tissue.

Quality control was performed within the HIDE-Deconv framework at both the cell and gene level. Cells were retained if they contained between 500 and 20,000 counts, a mitochondrial fraction below 5%, and a hemoglobin fraction below 5%. Cells with MALAT1 expression above the 99th percentile were removed. Genes expressed in fewer than 10 cells were discarded. Cell types represented by fewer than 100 cells were excluded, affecting only a small stromal-cell population.

Before preprocessing, the sampled dataset contained 606,065 cells and 17,764 genes. After preprocessing, the dataset comprised 518,874 cells from 126 patients and 16,348 genes. Cell annotations were provided using a three-level hierarchy consisting of 10 major, 17 minor and 32 sub-minor cell-type populations.

For benchmarking, the dataset was partitioned at the study level to reduce information leakage and provide a stringent assessment of generalization across independent datasets. The atlas comprised 11 studies, of which 6 were assigned to the training partition and 5 to the test partition. Consequently, all cells originating from a given study were assigned exclusively to either the training or test partition. Cell-type distributions for the training and test partitions at the major, minor and sub-minor annotation levels are summarized in Supplementary Tables S1-3. The corresponding hierarchy used throughout the benchmarking analyses is provided in Supplementary Table S4.

The training partition was used for feature selection, reference construction and model training within the HIDE-Deconv framework, whereas the test partition was used exclusively for generation of benchmark pseudobulk datasets (see Supplementary Methods Section S1.4.2). For all benchmarked methods, reference construction and model training were based solely on the training partition. For all LUAD application analyses, the complete dataset was used for reference construction and model training.

##### 1.2.2 PBMC single-cell reference dataset

As a single-cell reference for peripheral blood mononuclear cell deconvolution analyses, we used the Immune Health Atlas published by Gong et al. (4), which is distributed through the Allen Institute for Immunology Immune Health Atlas resource (5). The dataset was obtained from the Allen Institute for Immunology data portal (6).

To reduce computational requirements while preserving relative cell-type frequencies, 10% of cells were randomly sampled independently from each annotated cell population prior to preprocessing. Quality control was performed using the same filtering criteria as applied to the LUAD reference dataset. Cells were retained if they contained between 500 and 20,000 counts, a mitochondrial fraction below 5%, and a hemoglobin fraction below 5%. Cells with MALAT1 expression above the 99th percentile were removed. Genes expressed in fewer than 10 cells were discarded. Cell types represented by fewer than 100 cells before preprocessing were removed. Following quality control, a single erythrocyte remained in the dataset. Because mature erythrocytes are not considered part of the peripheral blood mononuclear cell (PBMC) fraction and their abundance was insufficient for meaningful reference-profile construction, this population was excluded from further analyses.

Before preprocessing, the sampled dataset contained 182,169 cells and 33,538 genes. After quality control, the final reference dataset comprised 155,894 cells from 108 donors and 21,857 genes.

Cell-type annotations provided by the original study were retained. For this work, we used the first two levels of the annotation hierarchy, resulting in 7 major and 25 minor cell-type populations. Cell-type distributions across both hierarchy levels are summarized in Supplementary Tables S5-6. The corresponding hierarchy is provided in Supplementary Table S7.

The PBMC reference dataset was used for feature selection, reference construction and model training in the PBMC deconvolution analyses of sepsis, COVID-19 and systemic lupus erythematosus cohorts.

#### 1.3 Cross-platform normalization between single-cell and bulk RNA-seq data

To account for systematic differences between single-cell and bulk RNA-sequencing platforms, we estimated gene-specific cross-platform correction factors. First, a subset of bulk RNA-seq samples was reserved exclusively for estimating the transfer coefficients. In parallel, pseudobulk samples were generated from the corresponding single-cell reference dataset using the HIDE-Deconv framework (see Supplementary Methods S1.4.2).

For each gene $g$, the median expression across the simulated pseudobulk samples and the selected real bulk samples was calculated. Medians were chosen instead of means to reduce sensitivity to outlier expression values. The domain-transfer factor $\alpha_g$ was estimated as

$$\alpha_g = \frac{\text{median}(Y_g^{sc})}{\text{median}(Y_g^{bulk})}. \tag{6}$$

where $\text{median}(Y_g^{sc})$ and $\text{median}(Y_g^{bulk})$ denote the median expression of gene $g$ across simulated and real bulk samples, respectively.

The remaining bulk samples were subsequently adjusted by multiplying each gene by its corresponding transfer factor prior to deconvolution. To enable cross-platform normalization for all $N$ bulk samples, a cross-validation strategy was employed. In each fold, $d$ samples were reserved for estimating $\alpha$, while the remaining samples were corrected and deconvolved.

The subset used for estimating $\alpha$ was iteratively shifted across the complete dataset using a sliding-window procedure, ensuring that normalization factors were estimated independently of the samples subsequently subjected to deconvolution.

#### 1.4 Benchmark

##### 1.4.1 Benchmark methods

For benchmark analyses, feature selection, reference construction and model training were performed exclusively on the training partition to prevent information leakage between training and benchmark data. For application analyses, reference profiles were generated using the complete corresponding reference dataset.

For DTD, HIDE and HIDE-Deconv, feature selection and reference construction followed the procedures described for the respective methods. Briefly, genes were ranked according to variance across cell-type reference profiles and the 5,000 highest-ranking genes were retained.

BayesPrism, EPIC, MuSiC, PREDE and Rectangle were evaluated using the complete set of available reference genes. No additional cell-type-specific subsampling was performed. Unless otherwise stated, default software settings and recommended workflows were used.

Whenever compatible with the requirements of a given method, reference profiles derived from the training partition were used to ensure a consistent benchmarking setup across methods. See Table 2 for further information.

##### 1.4.2 Pseudobulk benchmark generation

Benchmark pseudobulk datasets were generated exclusively from cells assigned to the independent test partition of the LUAD reference dataset. Prior to pseudobulk generation, single-cell expression profiles were library-size normalized to a scaling factor of $10^4$. The normalized data were subsequently used for feature selection, reference construction and pseudobulk generation.

For each pseudobulk sample, 100 cells were randomly selected and their normalized expression profiles were summed, resulting in bulk-like expression profiles with known cellular compositions. To assess variability in benchmark performance, ten independent benchmark datasets were generated, each consisting of 1,000 pseudobulk samples. All reported benchmark results represent summary statistics across these independent repetitions.

The resulting pseudobulk profiles were used directly for methods operating on normalized expression values. When individual methods required alternative input formats, method-specific input matrices were generated while preserving the same underlying samples and selected feature set.

#### 1.5 Downstream analyses

##### 1.5.1 Dimensionality reduction and clustering

Partial least-squares discriminant analysis (7) was performed on deconvolved cell-type compositions using disease stage or phenotypic group assignments as response variables. The first two latent components were used for visualization. Loadings were used to assess the contribution of individual cell types to group separation.

##### 1.5.2 Gene ranking and marker visualization

To investigate the molecular features learned by HIDE-Deconv, genes were ranked according to

$$s_i = g_i \cdot \mathrm{Var}(X_i), \tag{7}$$

where $g_i$ denotes the learned gene weight and $\mathrm{Var}(X_i)$ the variance of gene $i$ across the corresponding cell-type reference profiles. This score prioritizes genes that receive high weights during deconvolution while simultaneously exhibiting strong cell-type-specific variation. For visualization, genes were ranked independently for each hierarchy level and the highest-scoring genes were displayed in the marker heatmaps.

##### 1.5.3 Tumor microenvironment state identification

To identify distinct tumor microenvironment (TME) states, samples were clustered based on deconvolved sub-minor cell-type abundances using k-means clustering. HIDE-Deconv supports user-defined cluster numbers, allowing different biological resolutions to be explored depending on the application. Principal component analysis was subsequently used to visualize the resulting cluster structure.

##### 1.5.4 Differential gene expression and hallmark analysis

Differential gene expression analysis was performed on raw RNA-sequencing count data using the pyDeSeq2 interface implemented in HIDE-Deconv (8). Samples were grouped according to their k-means cluster assignments, and differential gene expression was assessed for all pairwise cluster comparisons. Genes with an absolute log2 fold change $> 1$ and an adjusted p-value $< 0.05$ were considered differentially expressed. Separate lists of upregulated and downregulated genes were generated for each pairwise comparison and subsequently analyzed using the cancer hallmark analysis platform described by Menyhart et al. (9).

#### 1.6 Supplementary Tables

##### 1.6.1 LUAD hierarchy and cell distributions

Table 1: Summary of benchmarked deconvolution methods and implementation settings.

| Method | Implementation | Method class | Hierarchy | Remarks |
|---|---|---|---|---|
| HIDE-Deconv | Python package v0.1.10 | Joint weighted least squares | Yes | 20,000 pseudobulk samples; 2,000 training iterations; feature selection (5,000 genes) |
| DTD | Python package Deconomix v1.0.0 | Weighted least squares | No | 20,000 pseudobulk samples; feature selection (5,000 genes) |
| HIDE | code via zenodo (doi: 10.5281/zenodo.14724906) | Hierarchical weighted least squares | Yes | Top-down hierarchical inference; 20,000 pseudobulk samples; feature selection (5,000 genes) |
| BayesPrism / InstaPrism | R package InstaPrism v0.1.6 | Bayesian inference | Native (2-level) | Full reference dataset |
| PREDE | R package v1.2.1 | Partial-reference deconvolution | No | Hidden component disabled; full reference dataset |
| EPIC | R package v1.1.7 | Constrained regression | No | Cell-size factors estimated from training data; full reference dataset |
| MuSiC | R package v1.0.0 | Weighted regression | No | Default weighted MuSiC estimator; full reference dataset |
| Rectangle | Python package v1.5.0 | Marker-based deconvolution | No | Full reference dataset |

| Major | Train (N) | Train (%) | Test (N) | Test (%) | Total (N) | Total (%) |
|---|---|---|---|---|---|---|
| B-cell | 8846 | 3.49 | 18714 | 5.35 | 27560 | 4.57 |
| NK-cell | 18121 | 7.15 | 23401 | 6.69 | 41522 | 6.88 |
| T-cell | 85260 | 33.64 | 138500 | 39.58 | 223760 | 37.08 |
| endothelial | 10666 | 4.21 | 7156 | 2.04 | 17822 | 2.95 |
| epithelial | 41375 | 16.32 | 29489 | 8.43 | 70864 | 11.74 |
| mast_cell | 4950 | 1.95 | 5942 | 1.70 | 10892 | 1.81 |
| myeloid | 75715 | 29.87 | 100035 | 28.59 | 175750 | 29.13 |
| neutrophil | 7 | 0.00 | 7475 | 2.14 | 7482 | 1.24 |
| plasma | 3383 | 1.33 | 15259 | 4.36 | 18642 | 3.09 |
| stromal | 5139 | 2.03 | 3983 | 1.14 | 9122 | 1.51 |
| Total | 253462 | 100.00 | 349954 | 100.00 | 603416 | 100.00 |

Table 2: Single Cell distribution of the LUAD dataset, both total and benchmark distribution, at major cell type annotation level.

| Minor | Train (N) | Train (%) | Test (N) | Test (%) | Total (N) | Total (%) |
|---|---|---|---|---|---|---|
| B-cell | 8846 | 3.49 | 18714 | 5.35 | 27560 | 4.57 |
| NK-cell | 18121 | 7.15 | 23401 | 6.69 | 41522 | 6.88 |
| T-cell | 85260 | 33.64 | 138500 | 39.58 | 223760 | 37.08 |
| airway | 17353 | 6.85 | 7321 | 2.09 | 24674 | 4.09 |
| alveolar | 8609 | 3.40 | 7855 | 2.24 | 16464 | 2.73 |
| dendritic | 8624 | 3.40 | 12119 | 3.46 | 20743 | 3.44 |
| fibroblast | 4117 | 1.62 | 2187 | 0.62 | 6304 | 1.04 |
| lymphatic | 1547 | 0.61 | 225 | 0.06 | 1772 | 0.29 |
| macrophage | 58974 | 23.27 | 61173 | 17.48 | 120147 | 19.91 |
| malignant | 15413 | 6.08 | 14313 | 4.09 | 29726 | 4.93 |
| mast_cell | 4950 | 1.95 | 5942 | 1.70 | 10892 | 1.81 |
| mesothelial | 231 | 0.09 | 58 | 0.02 | 289 | 0.05 |
| monocyte | 8117 | 3.20 | 26743 | 7.64 | 34860 | 5.78 |
| neutrophil | 7 | 0.00 | 7475 | 2.14 | 7482 | 1.24 |
| peri-vascular | 791 | 0.31 | 1738 | 0.50 | 2529 | 0.42 |
| plasma | 3383 | 1.33 | 15259 | 4.36 | 18642 | 3.09 |
| vascular | 9119 | 3.60 | 6931 | 1.98 | 16050 | 2.66 |
| Total | 253462 | 100.00 | 349954 | 100.00 | 603416 | 100.00 |

Table 3: Single Cell distribution of the LUAD dataset, both total and benchmark distribution, at minor cell type annotation level.

| Sub-Minor | Train (N) | Train (%) | Test (N) | Test (%) | Total (N) | Total (%) |
|---|---|---|---|---|---|---|
| B cell | 8846 | 3.49 | 18714 | 5.35 | 27560 | 4.57 |
| CD1c-positive myeloid dendritic cell | 6868 | 2.71 | 8507 | 2.43 | 15375 | 2.55 |
| CD4-positive, alpha-beta T cell | 36002 | 14.20 | 76834 | 21.96 | 112836 | 18.70 |
| CD8-positive, alpha-beta T cell | 40341 | 15.92 | 49604 | 14.17 | 89945 | 14.91 |
| alveolar macrophage | 34101 | 13.45 | 29518 | 8.43 | 63619 | 10.54 |
| bronchus fibroblast of lung | 1150 | 0.45 | 977 | 0.28 | 2127 | 0.35 |
| capillary endothelial cell | 2988 | 1.18 | 2253 | 0.64 | 5241 | 0.87 |
| classical monocyte | 6544 | 2.58 | 23182 | 6.62 | 29726 | 4.93 |
| club cell | 402 | 0.16 | 652 | 0.19 | 1054 | 0.17 |
| conventional dendritic cell | 610 | 0.24 | 1057 | 0.30 | 1667 | 0.28 |
| dendritic cell | 535 | 0.21 | 634 | 0.18 | 1169 | 0.19 |
| endothelial cell of lymphatic vessel | 1547 | 0.61 | 225 | 0.06 | 1772 | 0.29 |
| epithelial cell of lung | 13387 | 5.28 | 5328 | 1.52 | 18715 | 3.10 |
| fibroblast of lung | 2967 | 1.17 | 1210 | 0.35 | 4177 | 0.69 |
| macrophage | 22897 | 9.03 | 29478 | 8.42 | 52375 | 8.68 |
| malignant cell | 15413 | 6.08 | 14313 | 4.09 | 29726 | 4.93 |
| mast cell | 4950 | 1.95 | 5942 | 1.70 | 10892 | 1.81 |
| mesothelial cell | 231 | 0.09 | 58 | 0.02 | 289 | 0.05 |
| multiciliated epithelial cell | 3564 | 1.41 | 1341 | 0.38 | 4905 | 0.81 |
| myeloid cell | 1976 | 0.78 | 2177 | 0.62 | 4153 | 0.69 |
| natural killer cell | 18121 | 7.15 | 23401 | 6.69 | 41522 | 6.88 |
| neutrophil | 7 | 0.00 | 7475 | 2.14 | 7482 | 1.24 |
| non-classical monocyte | 1573 | 0.62 | 3561 | 1.02 | 5134 | 0.85 |
| pericyte | 428 | 0.17 | 786 | 0.22 | 1214 | 0.20 |
| plasma cell | 3383 | 1.33 | 15259 | 4.36 | 18642 | 3.09 |
| plasmacytoid dendritic cell | 611 | 0.24 | 1921 | 0.55 | 2532 | 0.42 |
| pulmonary alveolar type 1 cell | 2364 | 0.93 | 794 | 0.23 | 3158 | 0.52 |
| pulmonary alveolar type 2 cell | 6245 | 2.46 | 7061 | 2.02 | 13306 | 2.21 |
| pulmonary artery endothelial cell | 815 | 0.32 | 648 | 0.19 | 1463 | 0.24 |
| regulatory T cell | 8917 | 3.52 | 12062 | 3.45 | 20979 | 3.48 |
| smooth muscle cell | 363 | 0.14 | 952 | 0.27 | 1315 | 0.22 |
| vein endothelial cell | 5316 | 2.10 | 4030 | 1.15 | 9346 | 1.55 |
| Summe | 253462 | 100.00 | 349954 | 100.00 | 603416 | 100.00 |

Table 4: Single Cell distribution of the LUAD dataset, both total and benchmark distribution, at sub-minor cell type annotation level.

| Major | Minor | Sub |
|---|---|---|
| B-cell | B-cell | B cell |
| endothelial | lymphatic | endothelial cell of lymphatic vessel |
| endothelial | vascular | capillary endothelial cell |
| endothelial | vascular | pulmonary artery endothelial cell |
| endothelial | vascular | vein endothelial cell |
| epithelial | airway | club cell |
| epithelial | airway | epithelial cell of lung |
| epithelial | airway | multiciliated epithelial cell |
| epithelial | alveolar | pulmonary alveolar type 1 cell |
| epithelial | alveolar | pulmonary alveolar type 2 cell |
| epithelial | malignant | malignant cell |
| mast cell | mast cell | mast cell |
| myeloid | dendritic | CD1c-positive myeloid dendritic cell |
| myeloid | dendritic | conventional dendritic cell |
| myeloid | dendritic | dendritic cell |
| myeloid | dendritic | plasmacytoid dendritic cell |
| myeloid | macrophage | alveolar macrophage |
| myeloid | macrophage | macrophage |
| myeloid | macrophage | myeloid cell |
| myeloid | monocyte | classical monocyte |
| myeloid | monocyte | non-classical monocyte |
| neutrophil | neutrophil | neutrophil |
| NK-cell | NK-cell | natural killer cell |
| plasma | plasma | plasma cell |
| stromal | fibroblast | bronchus fibroblast of lung |
| stromal | fibroblast | fibroblast of lung |
| stromal | mesothelial cell | mesothelial cell |
| stromal | peri-vascular | pericyte |
| stromal | peri-vascular | smooth muscle cell |
| T-cell | T-cell | CD4-positive, alpha-beta T cell |
| T-cell | T-cell | CD8-positive, alpha-beta T cell |
| T-cell | T-cell | regulatory T cell |

Table 5: Cell-type hierarchy of the LUAD dataset.

### 2 Cell Type Distribution and hierarchy of PBMC Dataset

| Major cell-type | Absolute cell count | Percentage |
|---|---|---|
| T cell | 101921 | 65.38% |
| Monocyte | 27074 | 17.37% |
| B cell | 12582 | 8.07% |
| NK cell | 11891 | 7.63% |
| DC | 2075 | 1.33% |
| Platelet | 190 | 0.12% |
| Progenitor cell | 161 | 0.10% |
| Total | 155894 | 100.00% |

Table 6: Table showing number of single cells per major cell-type for PBMC dataset.

| Minor cell-type | Absolute cell count | Percentage |
|---|---|---|
| Naive CD4 T cell | 36897 | 23.67% |
| Memory CD4 T cell | 28764 | 18.45% |
| CD14 monocyte | 22400 | 14.37% |
| Memory CD8 T cell | 13652 | 8.76% |
| Naive CD8 T cell | 11104 | 7.12% |
| CD56dim NK cell | 10648 | 6.83% |
| Naive B cell | 6724 | 4.31% |
| Memory B cell | 3764 | 2.41% |
| gdT | 3715 | 2.38% |
| Treg | 3696 | 2.37% |
| CD16 monocyte | 3579 | 2.30% |
| MAIT | 3246 | 2.08% |
| cDC2 | 1336 | 0.86% |
| Intermediate monocyte | 1095 | 0.70% |
| Transitional B cell | 1089 | 0.70% |
| CD56bright NK cell | 1023 | 0.66% |
| Effector B cell | 822 | 0.53% |
| pDC | 739 | 0.47% |
| CD8aa | 456 | 0.29% |
| Proliferating NK cell | 220 | 0.14% |
| Proliferating T cell | 210 | 0.13% |
| Platelet | 190 | 0.12% |
| Plasma cell | 183 | 0.12% |
| DN T cell | 181 | 0.12% |
| Progenitor cell | 161 | 0.10% |
| Total | 155894 | 100.00% |

Table 7: Table showing number of single cells per minor cell-type for PBMC dataset.

| Major | Minor |
| --- | --- |
| Monocyte | CD14 monocyte |
| Monocyte | CD16 monocyte |
| NK cell | CD56bright NK cell |
| NK cell | CD56dim NK cell |
| T cell | CD8aa |
| T cell | DN T cell |
| B cell | Effector B cell |
| Monocyte | Intermediate monocyte |
| T cell | MAIT |
| B cell | Memory B cell |
| T cell | Memory CD4 T cell |
| T cell | Memory CD8 T cell |
| B cell | Naive B cell |
| T cell | Naive CD4 T cell |
| T cell | Naive CD8 T cell |
| B cell | Plasma cell |
| Platelet | Platelet |
| Progenitor cell | Progenitor cell |
| NK cell | Proliferating NK cell |
| T cell | Proliferating T cell |
| B cell | Transitional B cell |
| T cell | Treg |
| DC | cDC2 |
| T cell | gdT |
| DC | pDC |

Table 8: Cell-type hierarchy used for the PBMC dataset.

### 3 Bulk Datasets

Table 9: Bulk RNA-seq datasets used for HIDE-Deconv application analyses.

| Dataset | Disease | Samples | Comparison | Accession | Reference |
|---|---|---|---|---|---|
| SU2C-MARK | LUAD | 110 | Pre-treatment LUAD tumors treated with PD-(L)1 blockade | dbGaP: phs002822.v1.p1 | Ravi et al., Nat. Genet. (2023) |
| GSE112087 | SLE | 60 | 31 SLE vs 29 healthy controls | GEO: GSE112087 | Oon et al., Clin. Transl. Immunol. (2019); Figgett et al., Clin. Transl. Immunol. (2019) |
| GSE152418 | COVID-19 | 34 | 17 COVID-19 vs 17 healthy controls | GEO: GSE152418 | Arunachalam et al., Science (2020) |
| GSE279448 | Sepsis | 89 | 46 adult sepsis patients vs 43 healthy controls | GEO: GSE279448 | Ye et al. (2025) |

### 4 Results Benchmark

#### 4.1 Spearman Correlation

| SCC | BayesPrism | DTD | EPIC | HIDE |
|---|---|---|---|---|
| Sub-minor | $0.645 \pm 0.004$ | $0.647 \pm 0.006$ | $0.156 \pm 0.005$ | $0.599 \pm 0.003$ |
| Minor | $0.719 \pm 0.004$ | $0.745 \pm 0.007$ | $0.221 \pm 0.007$ | $0.749 \pm 0.004$ |
| Major (aggregated) | $0.811 \pm 0.003$ | $0.864 \pm 0.002$ | $0.266 \pm 0.012$ | $0.913 \pm 0.003$ |
| Major (direct) | - | $0.874 \pm 0.002$ | $0.634 \pm 0.006$ | - |
| Comparison aggregated / direct | 1 | $0.879 \pm 0.003$ | $0.354 \pm 0.012$ | 1 |

| SCC | HIDE-Deconv | MuSiC | PREDE | Rectangle |
|---|---|---|---|---|
| Sub | $0.704 \pm 0.006$ | $0.583 \pm 0.005$ | $0.339 \pm 0.004$ | $0.614 \pm 0.004$ |
| Minor | $0.792 \pm 0.006$ | $0.665 \pm 0.007$ | $0.500 \pm 0.005$ | $0.717 \pm 0.005$ |
| Major (aggregated) | $0.902 \pm 0.002$ | $0.736 \pm 0.005$ | $0.626 \pm 0.006$ | $0.811 \pm 0.005$ |
| Major (direct) | - | $0.560 \pm 0.009$ | $0.553 \pm 0.007$ | $0.8 \pm 0.005$ |
| Comparison aggregated / direct | 1 | $0.832 \pm 0.003$ | $0.741 \pm 0.005$ | $0.813 \pm 0.003$ |

Table 10: Spearman Correlation for the sub-minor, minor and major cell-type level. The major level is calculated in two ways: Summing up the results of the sub-minor level (major (aggregated)) and also direct only on major cell-type level (major (direct)). Additionally the results of both aggregated and independent approach are compared to each other to provide an overview about the differences between them. A dash (-) indicates that this method uses a hierarchical setup an has therefore no direct major calculation.

#### 4.2 Pearson Correlation

| PCC | BayesPrism | DTD | EPIC | HIDE |
|---|---|---|---|---|
| Sub | $0.725 \pm 0.005$ | $0.687 \pm 0.007$ | $0.059 \pm 0.007$ | $0.642 \pm 0.005$ |
| Minor | $0.794 \pm 0.005$ | $0.77 \pm 0.001$ | $0.089 \pm 0.011$ | $0.783 \pm 0.005$ |
| Major (aggregated) | $0.82 \pm 0.003$ | $0.87 \pm 0.002$ | $0.097 \pm 0.012$ | $0.914 \pm 0.002$ |
| Major (direct) | - | $0.88 \pm 0.003$ | - | |
| Comparison aggregated / direct | 1 | $0.888 \pm 0.004$ | $0.132 \pm 0.02$ | 1 |

| PCC | HIDE-Deconv | MuSiC | PREDE | Rectangle |
|---|---|---|---|---|
| Sub | $0.758 \pm 0.005$ | $0.612 \pm 0.007$ | $0.394 \pm 0.005$ | $0.685 \pm 0.005$ |
| Minor | $0.83 \pm 0.007$ | $0.671 \pm 0.007$ | $0.557 \pm 0.005$ | $0.78 \pm 0.005$ |
| Major (aggregated) | $0.906 \pm 0.001$ | $0.736 \pm 0.004$ | $0.647 \pm 0.006$ | $0.817 \pm 0.004$ |
| Major (direct) | - | $0.71 \pm 0.004$ | $0.567 \pm 0.009$ | $0.801 \pm 0.005$ |
| Comparison aggregated / direct | 1 | $0.832 \pm 0.004$ | $0.76 \pm 0.003$ | $0.813 \pm 0.003$ |

Table 11: Pearson Correlation for the sub-minor, minor and major cell-type level. The major level is calculated in two ways: Summing up the results of the sub-minor level (major (aggregated)) and also direct only on major cell-type level (major (direct)). Additionally the results of both aggregated and independent approach are compared to each other to provide an overview about the differences between them. A dash (-) indicates that this method uses a hierarchical setup an has therefore no direct major calculation.

##### 4.3 N-Mean Absolute Error

| NMAE | BayesPrism | DTD | EPIC | HIDE |
|---|---|---|---|---|
| Sub | $1.071 \pm 0.018$ | $1.261 \pm 0.029$ | $12.147 \pm 1.322$ | $1.169 \pm 0.031$ |
| Minor | $0.699 \pm 0.019$ | $0.725 \pm 0.013$ | $16.433 \pm 2.462$ | $0.788 \pm 0.05$ |
| Major (aggregated) | $0.562 \pm 0.004$ | $0.42 \pm 0.004$ | $2.326 \pm 0.043$ | $0.24 \pm 0.002$ |
| Major (direct) | - | $0.426 \pm 0.003$ | $1.093 \pm 0.043$ | - |
| Comparison aggregated / direct | 0 | $0.3 \pm 0.003$ | $1.121 \pm 0.004$ | 0 |

| NMAE | HIDE-Deconv | MuSiC | PREDE | Rectangle |
|---|---|---|---|---|
| Sub | $0.791 \pm 0.02$ | $0.2.053 \pm 0.032$ | $2.286 \pm 0.042$ | $0.793 \pm 0.01$ |
| Minor | $0.706 \pm 0.03$ | $1.577 \pm 0.035$ | $1.226 \pm 0.022$ | $0.595 \pm 0.011$ |
| Major (aggregated) | $0.307 \pm 0.002$ | $0.1.665 \pm 0.012$ | $0.969 \pm 0.008$ | $0.527 \pm 0.005$ |
| Major (direct) | - | $2.694 \pm 0.018$ | $1.573 \pm 0.013$ | $0.805 \pm 0.009$ |
| Comparison aggregated / direct | 0 | $0.492 \pm 0.005$ | $1.2 \pm 0.003$ | $0.393 \pm 0.007$ |

Table 12: Normalized Mean Absolute Error for the sub-minor, minor and major cell-type level. The major level is calculated in two ways: Summing up the results of the sub-minor level (major (aggregated)) and also direct only on major cell-type level (major (direct)). Additionally the results of both aggregated and independent approach are compared to each other to provide an overview about the differences between them. A dash (-) indicates that this method uses a hierarchical setup an has therefore no direct major calculation.

### 5 LUAD Study

#### 5.1 Patient demographics

| Patient characteristics ($n = 110$) | All patients, no. | % |
|---|---|---|
| Age (years), median (range) | 64 (40 - 89) | |
| Sex | | |
| Male | 67 | 41 |
| Female | 65 | 59 |
| Smoking status | | |
| Never | 16 | 14.5 |
| Former | 78 | 71 |
| Current | 16 | 14.5 |
| Smoking pack-years | | |
| 0 | 16 | 15 |
| 1-10 | 16 | 15 |
| 11-20 | 17 | 15 |
| 21-40 | 35 | 32 |
| >40 | 23 | 21 |
| Histology | | |
| Adenocarcinoma | 110 | 100 |
| Initial Tumor Stage | | |
| Stage 1 | 10 | 9 |
| Stage 2 | 9 | 8 |
| Stage 3 | 21 | 19 |
| Stage 4 | 65 | 59 |
| Agent PD1 category | | |
| PD(L)1 | 96 | 87 |
| PD(L)1 + CTLA4 | 9 | 8 |
| PD(L)1 + Other | 5 | 5 |
| Agent PD1 received | | |
| Nivolumab | 52 | 47 |
| Pembrolizumab | 35 | 32 |
| Nivolumab + Ipilimumab | 8 | 7 |
| Other | 15 | 14 |
| BOR | | |
| PD | 36 | 33 |
| PR | 30 | 27 |
| SD | 28 | 25 |
| CR | 8 | 7 |

Table 13: The used LUAD samples consisted of 110 patients treated with immune checkpoint blockade therapy.

#### 5.2 Early vs Late - Mann Whitney U Major

| celltype | diff [Late-Early] | Late (3+4) mean ± std | Early (1+2) mean ± std | p | p_adj |
|---|---|---|---|---|---|
| T-cell | 0.1264 | 0.2319 ± 0.1593 | 0.1055 ± 0.0986 | 0.0002 | 0.0009 |
| epithelial | -0.2917 | 0.3037 ± 0.2554 | 0.5954 ± 0.2678 | 0.0001 | 0.0009 |
| NK-cell | 0.0296 | 0.0512 ± 0.0471 | 0.0216 ± 0.0157 | 0.0042 | 0.0142 |
| B-cell | 0.0538 | 0.0713 ± 0.1387 | 0.0175 ± 0.0236 | 0.0257 | 0.0514 |
| endothelial | -0.0048 | 0.0181 ± 0.0186 | 0.0228 ± 0.0122 | 0.0244 | 0.0514 |
| myeloid | 0.0771 | 0.2712 ± 0.1711 | 0.1941 ± 0.1224 | 0.0769 | 0.1282 |
| neutrophil | 0.0069 | 0.0102 ± 0.0203 | 0.0033 ± 0.0041 | 0.1322 | 0.1652 |
| plasma | -0.0004 | 0.0175 ± 0.0216 | 0.0179 ± 0.0345 | 0.1317 | 0.1652 |
| stromal | 0.0021 | 0.0137 ± 0.0166 | 0.0115 ± 0.0169 | 0.3936 | 0.4373 |
| mast cellcell | 0.0009 | 0.0112 ± 0.0107 | 0.0104 ± 0.0135 | 0.4595 | 0.4595 |

Table 14: Mann Whitney U of early vs late stage at major cell type level

#### 5.3 Early vs Late - Mann Whitney U Minor

| celltype | diff [Late-Early] | Late (3+4) mean ± std | Early (1+2) mean ± std | p | p_adj |
|---|---|---|---|---|---|
| T-cell | 0.1264 | $0.2319 \pm 0.1593$ | $0.1055 \pm 0.0986$ | 0.0002 | 0.0022 |
| alveolar | -0.2519 | $0.1390 \pm 0.2200$ | $0.3909 \pm 0.2770$ | 0.0003 | 0.0022 |
| NK-cell | 0.0296 | $0.0512 \pm 0.0471$ | $0.0216 \pm 0.0157$ | 0.0042 | 0.0234 |
| monocyte | 0.0318 | $0.0660 \pm 0.0626$ | $0.0343 \pm 0.0401$ | 0.0055 | 0.0234 |
| B-cell | 0.0538 | $0.0713 \pm 0.1387$ | $0.0175 \pm 0.0236$ | 0.0257 | 0.0828 |
| peri-vascular | 0.0015 | $0.0021 \pm 0.0042$ | $0.0006 \pm 0.0021$ | 0.0292 | 0.0828 |
| vascular | -0.0047 | $0.0149 \pm 0.0149$ | $0.0196 \pm 0.0132$ | 0.0519 | 0.1262 |
| mesothelial | 0.0010 | $0.0011 \pm 0.0047$ | $5.59 \cdot 10^{-5} \pm 0.0002$ | 0.0695 | 0.1477 |
| plasma | -0.0004 | $0.0175 \pm 0.0216$ | $0.0179 \pm 0.0345$ | 0.1317 | 0.2247 |
| neutrophil | 0.0069 | $0.0102 \pm 0.0204$ | $0.0033 \pm 0.0041$ | 0.1322 | 0.2247 |
| airway | -0.0393 | $0.0732 \pm 0.1231$ | $0.1125 \pm 0.1951$ | 0.2389 | 0.3692 |
| macrophage | 0.0437 | $0.1842 \pm 0.1532$ | $0.1405 \pm 0.0998$ | 0.3119 | 0.4418 |
| mast cellcell | 0.0009 | $0.0112 \pm 0.0107$ | $0.0104 \pm 0.0135$ | 0.4595 | 0.6009 |
| malignant | -0.0005 | $0.0915 \pm 0.0853$ | $0.0920 \pm 0.0611$ | 0.5480 | 0.6654 |
| dendritic | 0.0016 | $0.0210 \pm 0.0225$ | $0.0194 \pm 0.0159$ | 0.6987 | 0.7919 |
| lymphatic | $-4.41 \cdot 10^{-5}$ | $0.0032 \pm 0.0077$ | $0.0032 \pm 0.0056$ | 0.9813 | 0.9813 |
| fibroblast | -0.0004 | $0.0104 \pm 0.0134$ | $0.0108 \pm 0.0154$ | 0.9370 | 0.9813 |

Table 15: Mann Whitney U of early vs late stage at minor cell type level

#### 5.4 Early vs Late - Mann Whitney U Sub-Minor

| celltype | diff [Late-Early] | Late (3+4) mean ± std | Early (1+2) mean ± std | p | p_adj |
|---|---|---|---|---|---|
| pulmonary alveolar type 2 cell | -0.2559 | $0.1332 \pm 0.2194$ | $0.3891 \pm 0.2786$ | $9.97 \cdot 10^{-5}$ | 0.0032 |
| club cell | -0.0334 | $0.01604 \pm 0.0376$ | $0.0494 \pm 0.0757$ | 0.0011 | 0.0180 |
| vein endothelial cell | 0.0040 | $0.0075 \pm 0.0130$ | $0.0035 \pm 0.0087$ | 0.0053 | 0.0212 |
| capillary endothelial cell | -0.0100 | $0.0052 \pm 0.0080$ | $0.0152 \pm 0.0149$ | 0.0027 | 0.0212 |
| classical monocyte | 0.0345 | $0.0602 \pm 0.0643$ | $0.0257 \pm 0.0438$ | 0.0052 | 0.0212 |
| natural killer cell | 0.0296 | $0.0512 \pm 0.0471$ | $0.0216 \pm 0.0157$ | 0.0042 | 0.0212 |
| dendritic cell | 0.0022 | $0.0023 \pm 0.0047$ | $4.15 \cdot 10^{-5} \pm 0.0002$ | 0.0053 | 0.0212 |
| myeloid cell | 0.0140 | $0.0203 \pm 0.0329$ | $0.0062 \pm 0.0188$ | 0.0051 | 0.0212 |
| pulmonary alveolar type 1 cell | 0.0040 | $0.0057 \pm 0.0128$ | $0.0018 \pm 0.0041$ | 0.0061 | 0.0230 |
| regulatory T cell | 0.0186 | $0.0330 \pm 0.0345$ | $0.0144 \pm 0.0196$ | 0.0121 | 0.0369 |
| CD8-positive, alpha-beta T cell | 0.0761 | $0.1305 \pm 0.1361$ | $0.0544 \pm 0.0683$ | 0.0127 | 0.0369 |
| non-classical monocyte | -0.0028 | $0.0058 \pm 0.0089$ | $0.0086 \pm 0.0070$ | 0.0275 | 0.0678 |
| B cell | 0.0538 | $0.0713 \pm 0.1387$ | $0.0175 \pm 0.0236$ | 0.0257 | 0.0678 |
| smooth muscle cell | 0.0011 | $0.0011 \pm 0.0034$ | $5.28 \cdot 10^{-5} \pm 0.0001$ | 0.0310 | 0.0707 |
| mesothelial cell | 0.0010 | $0.0011 \pm 0.0047$ | $5.59 \cdot 10^{-5} \pm 0.0002$ | 0.0695 | 0.1390 |
| conventional dendritic cell | -0.0028 | $0.0027 \pm 0.0037$ | $0.0055 \pm 0.0080$ | 0.0662 | 0.1390 |
| macrophage | 0.0197 | $0.0384 \pm 0.0690$ | $0.0188 \pm 0.0455$ | 0.0871 | 0.1544 |
| pulmonary artery endothelial cell | 0.0013 | $0.0022 \pm 0.0049$ | $0.0009 \pm 0.0019$ | 0.0825 | 0.1544 |
| pericyte | 0.0004 | $0.0010 \pm 0.0023$ | $0.0006 \pm 0.0021$ | 0.0917 | 0.1544 |
| neutrophil | 0.0069 | $0.0102 \pm 0.0204$ | $0.0033 \pm 0.0041$ | 0.1322 | 0.2014 |
| plasma cell | -0.0004 | $0.0175 \pm 0.0216$ | $0.0179 \pm 0.0345$ | 0.1317 | 0.2014 |
| plasmacytoid dendritic cell | $-2.26 \cdot 10^{-6}$ | $0.0028 \pm 0.0041$ | $0.0028 \pm 0.0021$ | 0.1391 | 0.2023 |
| CD1c-positive myeloid dendritic cell | 0.0022 | $0.0133 \pm 0.0219$ | $0.0111 \pm 0.0124$ | 0.1543 | 0.2107 |
| multiciliated epithelial cell | 0.0108 | $0.0230 \pm 0.0717$ | $0.0122 \pm 0.0169$ | 0.1581 | 0.2107 |
| alveolar macrophage | 0.0100 | $0.1255 \pm 0.1426$ | $0.1155 \pm 0.0775$ | 0.2356 | 0.3015 |
| CD4-positive, alpha-beta T cell | 0.0318 | $0.0685 \pm 0.0866$ | $0.0367 \pm 0.0613$ | 0.2518 | 0.3099 |
| epithelial cell of lung | -0.0167 | $0.0342 \pm 0.0909$ | $0.0509 \pm 0.1867$ | 0.3189 | 0.3780 |
| bronchus fibroblast of lung | -0.0016 | $0.0045 \pm 0.0063$ | $0.0061 \pm 0.0081$ | 0.3369 | 0.3850 |
| fibroblast of lung | 0.0012 | $0.0059 \pm 0.0105$ | $0.0047 \pm 0.0083$ | 0.3811 | 0.4205 |
| mast cell | 0.0009 | $0.0112 \pm 0.0107$ | $0.0104 \pm 0.0135$ | 0.4595 | 0.4902 |
| malignant cell | -0.0005 | $0.0915 \pm 0.0853$ | $0.0920 \pm 0.0611$ | 0.5480 | 0.5656 |
| endothelial cell of lymphatic vessel | $-4.41 \cdot 10^{-5}$ | $0.0032 \pm 0.0077$ | $0.0032 \pm 0.0056$ | 0.9813 | 0.9813 |

Table 16: Mann Whitney U of early vs late stage at sub-cellular cell type level

#### 5.5 Early vs Late - Heatmap Visualization

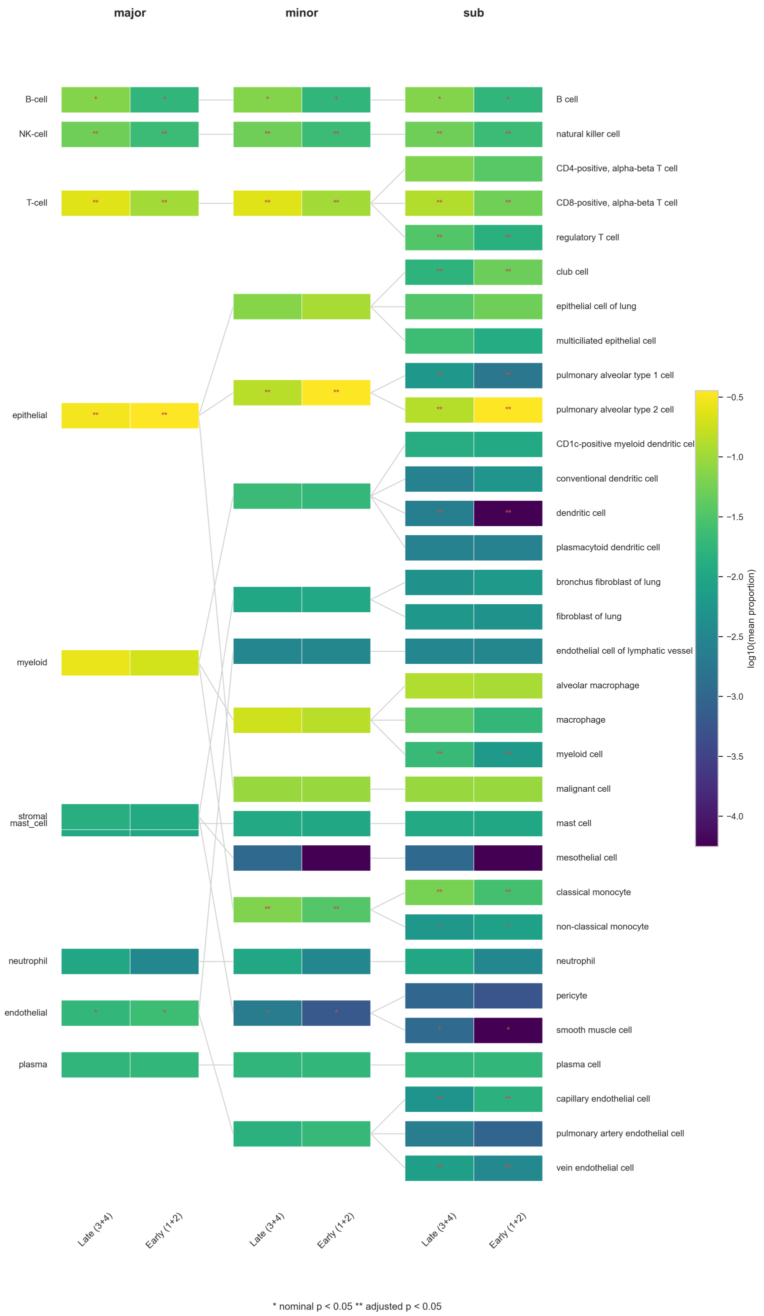

Figure 1: Heatmap visualization of log10 mean cellular proportions at all three cell hierarchy level for the early and late LUAD cohort. Differences between cell types were evaluated with a Mann-Whitney-U test. Significance is highlighted by either "**" for adjusted p-values below 0.05 or "*" for nominal p-values below 0.05.

##### 5.6 LUAD Markerplot Sub

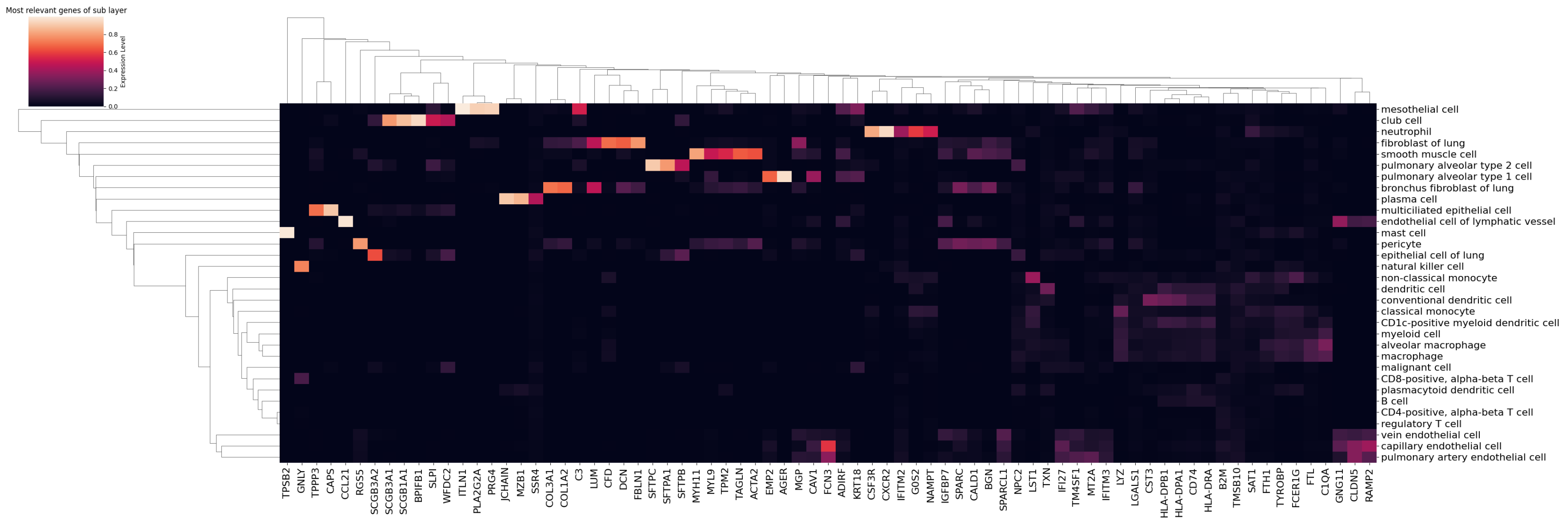

Figure 2: Sub cell type signatures learned by HIDE-Deconv. Shown are the 75 highest-ranked genes according to the score $s_i = g_i \cdot \mathrm{Var}(X_i)$, which combines learned gene weights with expression variability. Gene expression values were scaled between 0 and 1 for visualization.

#### 5.7 LUAD K-Mean PCA plots

### 5.7.1 k=2

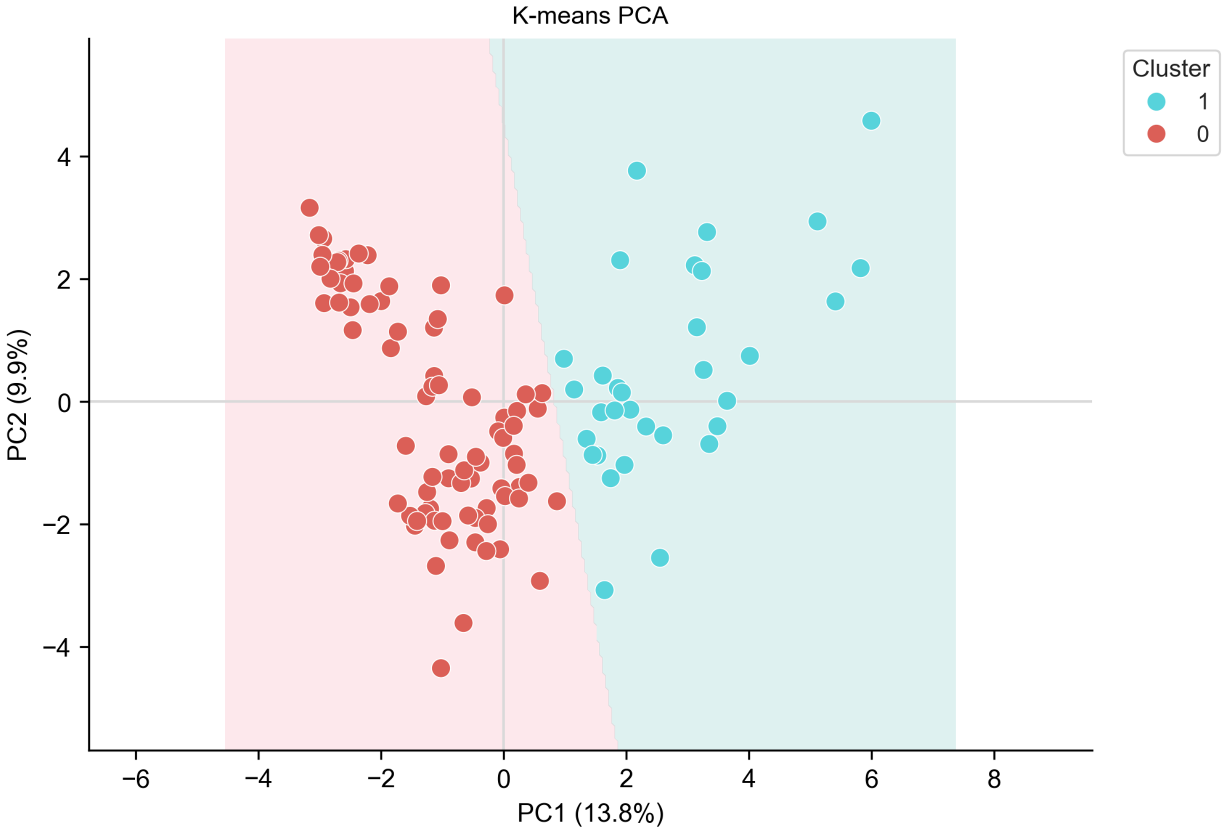

Figure 3: PCA plot of LUAD patient cluster assignment using $k = 2$ in HIDE-deconvs k-mean module.

#### 5.7.2 k=3

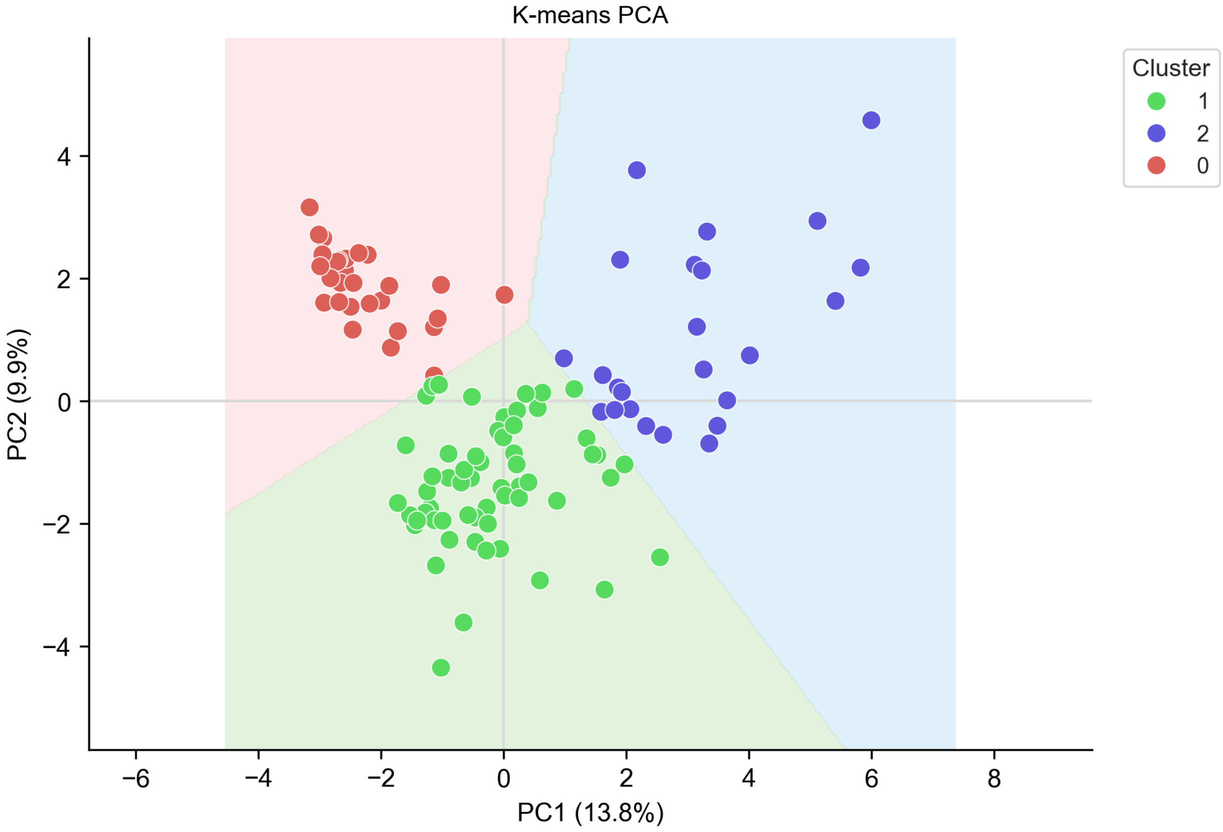

Figure 4: PCA plot of LUAD patient cluster assignment using $k = 3$ in HIDE-deconvs k-mean module.

### 5.7.3 k=4

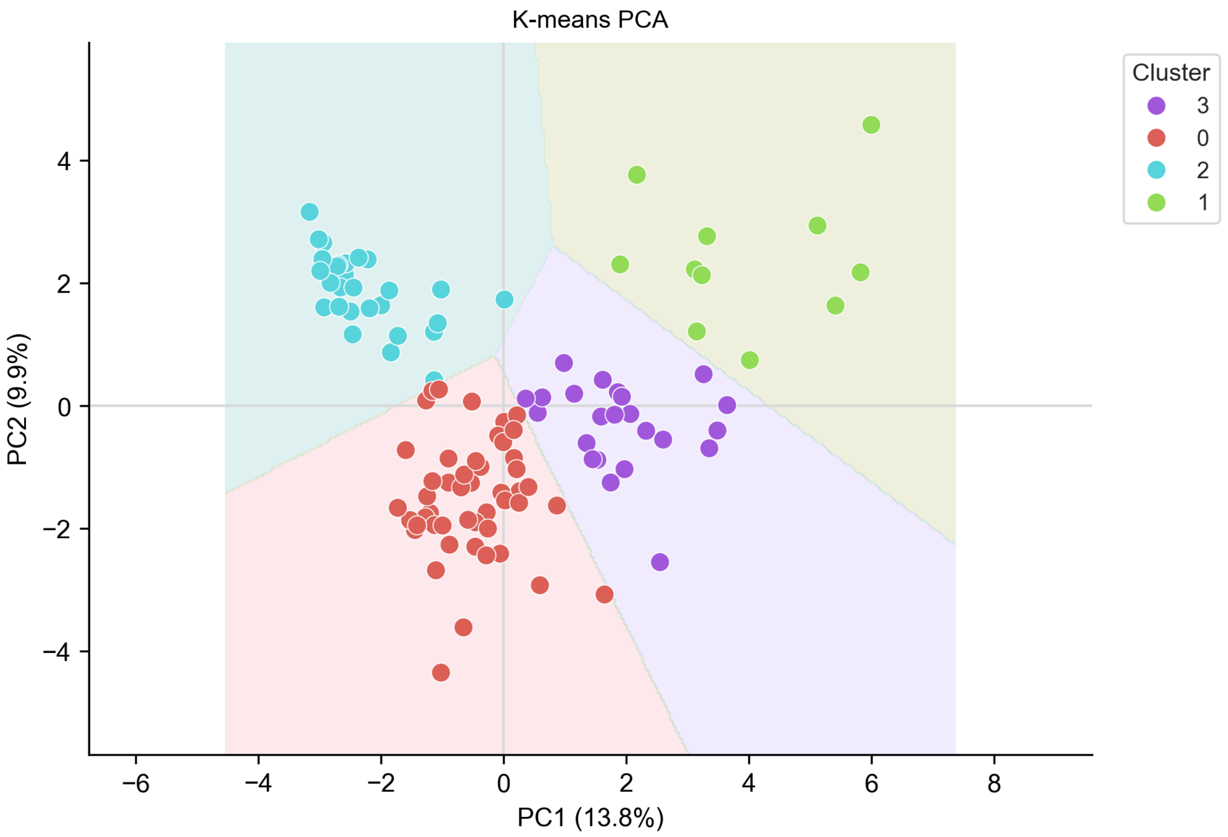

Figure 5: PCA plot of LUAD patient cluster assignment using $k = 4$ in HIDE-deconvs k-mean module.

#### 5.8 LUAD Clustering - Posthoc Dunn Major

| celltype | cohort 1 | cohort 2 | p_adj | mean [cohort 1] | mean [cohort 2] | diff [1-2] |
|---|---|---|---|---|---|---|
| epithelial | 0 | 1 | $2.78 \cdot 10^{-13}$ | 0.7371 | 0.2160 | 0.5211 |
| stromal | 1 | 2 | $1.07 \cdot 10^{-10}$ | 0.0073 | 0.0368 | -0.0295 |
| myeloid | 0 | 1 | $1.71 \cdot 10^{-9}$ | 0.1078 | 0.3112 | -0.2034 |
| T-cell | 0 | 1 | $2.51 \cdot 10^{-9}$ | 0.0797 | 0.2736 | -0.1939 |
| myeloid | 0 | 2 | $1.94 \cdot 10^{-8}$ | 0.1078 | 0.3161 | -0.2083 |
| epithelial | 0 | 2 | $3.99 \cdot 10^{-8}$ | 0.7371 | 0.2193 | 0.5178 |
| endothelial | 1 | 2 | $1.16 \cdot 10^{-7}$ | 0.0112 | 0.0354 | -0.0242 |
| stromal | 0 | 2 | $1.28 \cdot 10^{-7}$ | 0.0061 | 0.0368 | -0.0307 |
| mast cell | 1 | 2 | $2.37 \cdot 10^{-6}$ | 0.0078 | 0.0232 | -0.0154 |
| NK-cell | 0 | 2 | $4.03 \cdot 10^{-6}$ | 0.0196 | 0.0676 | -0.0480 |
| T-cell | 0 | 2 | $7.26 \cdot 10^{-6}$ | 0.0797 | 0.2234 | -0.1437 |
| mast cell | 0 | 2 | $1.13 \cdot 10^{-5}$ | 0.0072 | 0.0232 | -0.0160 |
| plasma | 0 | 2 | $4.92 \cdot 10^{-5}$ | 0.0050 | 0.0273 | -0.0223 |
| neutrophil | 0 | 1 | $5.70 \cdot 10^{-5}$ | 0.0013 | 0.0126 | -0.0113 |
| NK-cell | 0 | 1 | 0.0011 | 0.0196 | 0.0504 | -0.0308 |
| endothelial | 0 | 1 | 0.0013 | 0.0199 | 0.0112 | 0.0088 |
| plasma | 0 | 1 | 0.0042 | 0.0050 | 0.0195 | -0.0145 |
| neutrophil | 0 | 2 | 0.0154 | 0.0013 | 0.0093 | -0.0080 |
| NK-cell | 1 | 2 | 0.0194 | 0.0504 | 0.0676 | -0.0172 |
| plasma | 1 | 2 | 0.0352 | 0.0195 | 0.0273 | -0.0078 |
| endothelial | 0 | 2 | 0.0416 | 0.0199 | 0.0354 | -0.0155 |
| B-cell | 0 | 1 | 0.0416 | 0.0163 | 0.0904 | -0.0741 |
| B-cell | 0 | 2 | 0.0846 | 0.0163 | 0.0416 | -0.0253 |
| neutrophil | 1 | 2 | 0.2657 | 0.0126 | 0.0093 | 0.0032 |
| epithelial | 1 | 2 | 0.4907 | 0.2160 | 0.2193 | -0.0033 |
| myeloid | 1 | 2 | 0.5219 | 0.3112 | 0.3161 | -0.0049 |
| T-cell | 1 | 2 | 0.5620 | 0.2736 | 0.2234 | 0.0502 |
| stromal | 0 | 1 | 0.6099 | 0.0061 | 0.0073 | -0.0012 |
| mast cell | 0 | 1 | 0.8378 | 0.0072 | 0.0078 | -0.0006 |
| B-cell | 1 | 2 | 0.8840 | 0.0904 | 0.0416 | 0.0488 |

Table 17: Posthoc Dunn Test of clustered LUAD patients at major cell type level

#### 5.9 LUAD Clustering - Posthoc Dunn Minor

| celltype | cohort 1 | cohort 2 | p_adj | mean [cohort 1] | mean [cohort 2] | diff [1-2] |
|---|---|---|---|---|---|---|
| alveolar | 0 | 1 | $2.66 \cdot 10^{-14}$ | 0.5696 | 0.0502 | 0.5194 |
| peri-vascular | 0 | 2 | $1.76 \cdot 10^{-12}$ | 0.0002 | 0.0066 | -0.0064 |
| fibroblast | 1 | 2 | $1.26 \cdot 10^{-10}$ | 0.0051 | 0.0296 | -0.0245 |
| peri-vascular | 1 | 2 | $9.53 \cdot 10^{-10}$ | 0.0006 | 0.0066 | -0.0060 |
| T-cell | 0 | 1 | $2.51 \cdot 10^{-9}$ | 0.0797 | 0.2736 | -0.1939 |
| lymphatic | 0 | 2 | $9.49 \cdot 10^{-9}$ | 0.0002 | 0.0073 | -0.0071 |
| monocyte | 0 | 1 | $4.75 \cdot 10^{-8}$ | 0.0203 | 0.0824 | -0.0621 |
| alveolar | 0 | 2 | $4.87 \cdot 10^{-8}$ | 0.5696 | 0.0327 | 0.5369 |
| vascular | 1 | 2 | $4.99 \cdot 10^{-8}$ | 0.0082 | 0.0281 | -0.0199 |
| macrophage | 0 | 2 | $1.64 \cdot 10^{-7}$ | 0.0734 | 0.2314 | -0.1580 |
| macrophage | 0 | 1 | $4.25 \cdot 10^{-7}$ | 0.0734 | 0.2097 | -0.1363 |
| lymphatic | 1 | 2 | $1.85 \cdot 10^{-6}$ | 0.0029 | 0.0073 | -0.0044 |
| mast cell | 1 | 2 | $2.37 \cdot 10^{-6}$ | 0.0078 | 0.0232 | -0.0154 |
| vascular | 0 | 1 | $3.45 \cdot 10^{-6}$ | 0.0198 | 0.0082 | 0.0116 |
| NK-cell | 0 | 2 | $4.03 \cdot 10^{-6}$ | 0.0196 | 0.0676 | -0.0480 |
| T-cell | 0 | 2 | $7.26 \cdot 10^{-6}$ | 0.0797 | 0.2234 | -0.1437 |
| mast cell | 0 | 2 | $1.13 \cdot 10^{-5}$ | 0.0072 | 0.0232 | -0.0160 |
| fibroblast | 0 | 2 | $2.41 \cdot 10^{-5}$ | 0.0059 | 0.0296 | -0.0237 |
| plasma | 0 | 2 | $4.92 \cdot 10^{-5}$ | 0.0050 | 0.0273 | -0.0223 |
| neutrophil | 0 | 1 | $5.70 \cdot 10^{-5}$ | 0.0013 | 0.0126 | -0.0113 |
| monocyte | 0 | 2 | 0.0002 | 0.0203 | 0.0516 | -0.0313 |
| mesothelial | 0 | 2 | 0.0008 | $7.18 \cdot 10^{-6}$ | 0.0006 | -0.0006 |
| NK-cell | 0 | 1 | 0.0011 | 0.0196 | 0.0504 | -0.0308 |
| mesothelial | 0 | 1 | 0.0026 | $7.18 \cdot 10^{-6}$ | 0.0016 | -0.0016 |
| malignant | 0 | 1 | 0.0037 | 0.1160 | 0.0752 | 0.0407 |
| plasma | 0 | 1 | 0.0042 | 0.0050 | 0.0195 | -0.0145 |
| dendritic | 1 | 2 | 0.0047 | 0.0191 | 0.0332 | -0.0141 |
| dendritic | 0 | 2 | 0.0100 | 0.0141 | 0.0332 | -0.0191 |
| neutrophil | 0 | 2 | 0.0154 | 0.0013 | 0.0093 | -0.0080 |
| NK-cell | 1 | 2 | 0.0194 | 0.0504 | 0.0676 | -0.0172 |
| peri-vascular | 0 | 1 | 0.0287 | 0.0002 | 0.0006 | -0.0005 |
| plasma | 1 | 2 | 0.0352 | 0.0195 | 0.0273 | -0.0078 |
| lymphatic | 0 | 1 | 0.0409 | 0.0001 | 0.0029 | -0.0028 |
| B-cell | 0 | 1 | 0.0416 | 0.0163 | 0.0904 | -0.0741 |
| fibroblast | 0 | 1 | 0.0814 | 0.0059 | 0.0051 | 0.0008 |
| B-cell | 0 | 2 | 0.0846 | 0.0163 | 0.0416 | -0.0253 |
| malignant | 1 | 2 | 0.0859 | 0.0752 | 0.1034 | -0.0282 |
| macrophage | 1 | 2 | 0.1734 | 0.2097 | 0.2314 | -0.0217 |
| mesothelial | 1 | 2 | 0.2276 | 0.0016 | 0.0006 | 0.0010 |
| neutrophil | 1 | 2 | 0.2657 | 0.0126 | 0.0093 | 0.0032 |
| alveolar | 1 | 2 | 0.3092 | 0.0502 | 0.0327 | 0.0175 |
| vascular | 0 | 2 | 0.3098 | 0.0198 | 0.0281 | -0.0083 |
| monocyte | 1 | 2 | 0.3100 | 0.0824 | 0.0516 | 0.0309 |
| malignant | 0 | 2 | 0.3108 | 0.1160 | 0.1034 | 0.0126 |
| T-cell | 1 | 2 | 0.5620 | 0.2736 | 0.2234 | 0.0502 |
| mast cell | 0 | 1 | 0.8378 | 0.0072 | 0.0078 | -0.0006 |
| B-cell | 1 | 2 | 0.8840 | 0.0904 | 0.0416 | 0.0488 |
| dendritic | 0 | 1 | 0.9552 | 0.0141 | 0.0191 | -0.0050 |

Table 18: Posthoc Dunn Test of clustered LUAD patients at minor cell type level

#### 5.10 LUAD Clustering - Posthoc Dunn Sub-Minor

| celltype | cohort_1 | cohort_2 | p_adj | mean [cohort_1] | mean [cohort_2] | diff [1-2] |
|---|---|---|---|---|---|---|
| pulmonary alveolar type 2 cell | 0 | 1 | $1.11 \cdot 10^{-13}$ | 0.5653 | 0.0464 | 0.5189 |
| capillary endothelial cell | 0 | 1 | $1.83 \cdot 10^{-10}$ | 0.0182 | 0.0023 | 0.0159 |
| vein endothelial cell | 0 | 2 | $6.42 \cdot 10^{-10}$ | 0.0012 | 0.0224 | -0.0212 |
| classical monocyte | 0 | 1 | $2.17 \cdot 10^{-9}$ | 0.0085 | 0.0769 | -0.0684 |
| pulmonary alveolar type 2 cell | 0 | 2 | $5.66 \cdot 10^{-9}$ | 0.5653 | 0.0236 | 0.5417 |
| endothelial cell of lymphatic vessel | 0 | 2 | $9.49 \cdot 10^{-9}$ | 0.0002 | 0.0073 | -0.0071 |
| fibroblast of lung | 0 | 2 | $2.62 \cdot 10^{-8}$ | 0.0018 | 0.0179 | -0.0161 |
| conventional dendritic cell | 0 | 1 | $3.25 \cdot 10^{-8}$ | 0.0047 | 0.0011 | 0.0037 |
| CD4-positive, alpha-beta T cell | 0 | 2 | $3.44 \cdot 10^{-8}$ | 0.0133 | 0.1297 | -0.1164 |
| fibroblast of lung | 1 | 2 | $5.84 \cdot 10^{-8}$ | 0.0026 | 0.0179 | -0.0153 |
| myeloid cell | 0 | 2 | $1.08 \cdot 10^{-7}$ | 0.0025 | 0.0349 | -0.0324 |
| pulmonary alveolar type 1 cell | 0 | 2 | $1.62 \cdot 10^{-7}$ | 0.0043 | 0.0091 | -0.0048 |
| vein endothelial cell | 1 | 2 | $1.74 \cdot 10^{-7}$ | 0.0033 | 0.0224 | -0.0191 |
| smooth muscle cell | 0 | 2 | $4.00 \cdot 10^{-7}$ | $1.30 \cdot 10^{-5}$ | 0.0033 | -0.0033 |
| pericyte | 0 | 2 | $4.76 \cdot 10^{-7}$ | 0.0001 | 0.0033 | -0.0031 |
| non-classical monocyte | 0 | 2 | $7.58 \cdot 10^{-7}$ | 0.0118 | 0.0027 | 0.0091 |
| non-classical monocyte | 0 | 1 | $8.49 \cdot 10^{-7}$ | 0.0118 | 0.0055 | 0.0063 |
| pericyte | 1 | 2 | $1.09 \cdot 10^{-9}$ | 0.0003 | 0.0033 | -0.0030 |
| endothelial cell of lymphatic vessel | 1 | 2 | $1.85 \cdot 10^{-6}$ | 0.0029 | 0.0073 | -0.0044 |
| capillary endothelial cell | 0 | 2 | $2.20 \cdot 10^{-6}$ | 0.0182 | 0.0040 | 0.0142 |
| mast cell | 1 | 2 | $2.37 \cdot 10^{-6}$ | 0.0078 | 0.0232 | -0.0154 |
| natural killer cell | 0 | 2 | $4.03 \cdot 10^{-6}$ | 0.0196 | 0.0676 | -0.0480 |
| club cell | 0 | 1 | $7.88 \cdot 10^{-6}$ | 0.0328 | 0.0133 | 0.0196 |
| mast cell | 0 | 2 | $1.13 \cdot 10^{-5}$ | 0.0072 | 0.0232 | -0.0160 |
| pulmonary artery endothelial cell | 0 | 1 | $1.33 \cdot 10^{-5}$ | 0.0004 | 0.0027 | -0.0023 |
| classical monocyte | 0 | 2 | $1.59 \cdot 10^{-5}$ | 0.0085 | 0.0488 | -0.0403 |
| conventional dendritic cell | 1 | 2 | $1.69 \cdot 10^{-5}$ | 0.0011 | 0.0061 | -0.0050 |
| dendritic cell | 0 | 1 | $2.03 \cdot 10^{-5}$ | 0.0 | 0.0029 | -0.0029 |
| pulmonary alveolar type 1 cell | 0 | 1 | $2.48 \cdot 10^{-5}$ | 0.0043 | 0.0038 | 0.0005 |
| epithelial cell of lung | 0 | 1 | $2.58 \cdot 10^{-5}$ | 0.0041 | 0.0549 | -0.0508 |

| | | | | | | |
|---|---|---|---|---|---|---|
| CD4-positive, alpha-beta T cell | 1 | 2 | $2.84 \cdot 10^{-5}$ | 0.0626 | 0.1297 | -0.0671 |
| bronchus fibroblast of lung | 1 | 2 | $4.30 \cdot 10^{-5}$ | 0.0026 | 0.0117 | -0.0091 |
| plasma cell | 0 | 2 | $4.92 \cdot 10^{-5}$ | 0.0050 | 0.0273 | -0.0223 |
| CD8-positive, alpha-beta T cell | 0 | 1 | $5.58 \cdot 10^{-5}$ | 0.0488 | 0.1719 | -0.1231 |
| neutrophil | 0 | 1 | $5.70 \cdot 10^{-5}$ | 0.0013 | 0.0126 | -0.0113 |
| smooth muscle cell | 1 | 2 | 0.0002 | 0.0003 | 0.0033 | -0.0030 |
| epithelial cell of lung | 0 | 2 | 0.0003 | 0.0041 | 0.0347 | -0.0307 |
| myeloid cell | 0 | 1 | 0.0006 | 0.0025 | 0.0180 | -0.0155 |
| mesothelial cell | 0 | 2 | 0.0008 | $7.18 \cdot 10^{-6}$ | 0.0006 | -0.0006 |
| natural killer cell | 0 | 1 | 0.0011 | 0.0196 | 0.0504 | -0.0308 |
| macrophage | 0 | 2 | 0.0015 | 0.0057 | 0.0583 | -0.0526 |
| mesothelial cell | 0 | 1 | 0.0026 | $7.18 \cdot 10^{-6}$ | 0.0016 | -0.0016 |
| myeloid cell | 1 | 2 | 0.0030 | 0.0180 | 0.0349 | -0.0170 |
| bronchus fibroblast of lung | 0 | 1 | 0.0035 | 0.0041 | 0.0026 | 0.0016 |
| malignant cell | 0 | 1 | 0.0037 | 0.1160 | 0.0752 | 0.0407 |
| CD8-positive, alpha-beta T cell | 1 | 2 | 0.0040 | 0.1719 | 0.0747 | 0.0972 |
| plasma cell | 0 | 1 | 0.0042 | 0.0050 | 0.0195 | -0.0145 |
| plasmacytoid dendritic cell | 1 | 2 | 0.0052 | 0.0019 | 0.0048 | -0.0029 |
| alveolar macrophage | 0 | 2 | 0.0055 | 0.0653 | 0.1382 | -0.0729 |
| alveolar macrophage | 0 | 1 | 0.0055 | 0.0653 | 0.1483 | -0.0830 |
| plasmacytoid dendritic cell | 0 | 1 | 0.0088 | 0.0025 | 0.0019 | 0.0006 |
| macrophage | 0 | 1 | 0.0090 | 0.0057 | 0.0434 | -0.0377 |
| pulmonary artery endothelial cell | 0 | 2 | 0.0102 | 0.0004 | 0.0018 | -0.0014 |
| dendritic cell | 0 | 2 | 0.0152 | 0.0 | 0.0020 | -0.0020 |
| neutrophil | 0 | 2 | 0.0154 | 0.0013 | 0.0093 | -0.0080 |
| club cell | 0 | 2 | 0.0157 | 0.0328 | 0.0271 | 0.0057 |
| CD4-positive, alpha-beta T cell | 0 | 1 | 0.0169 | 0.0133 | 0.0626 | -0.0493 |
| smooth muscle cell | 0 | 1 | 0.0193 | $1.30 \cdot 10^{-5}$ | 0.0003 | -0.0003 |
| natural killer cell | 1 | 2 | 0.0194 | 0.0504 | 0.0676 | -0.0172 |
| pulmonary alveolar type 1 cell | 1 | 2 | 0.0317 | 0.0038 | 0.0091 | -0.0053 |
| regulatory T cell | 1 | 2 | 0.0339 | 0.0391 | 0.0190 | 0.0202 |
| plasma cell | 1 | 2 | 0.0352 | 0.0195 | 0.0273 | -0.0078 |
| vein endothelial cell | 0 | 1 | 0.0369 | 0.0012 | 0.0033 | -0.0021 |
| endothelial cell of lymphatic vessel | 0 | 1 | 0.0409 | 0.0002 | 0.0029 | -0.0028 |
| B cell | 0 | 1 | 0.0416 | 0.0163 | 0.0904 | -0.0741 |
| B cell | 0 | 2 | 0.0846 | 0.0163 | 0.0416 | -0.0253 |
| malignant cell | 1 | 2 | 0.0859 | 0.0752 | 0.1034 | -0.0282 |
| regulatory T cell | 0 | 1 | 0.1016 | 0.0176 | 0.0391 | -0.0215 |
| club cell | 1 | 2 | 0.1287 | 0.0133 | 0.0271 | -0.0138 |
| macrophage | 1 | 2 | 0.1686 | 0.0434 | 0.0583 | -0.0149 |

| | | | | | | |
|---|---|---|---|---|---|---|
| dendritic cell | 1 | 2 | 0.1867 | 0.0029 | 0.0020 | 0.0008 |
| bronchus fibroblast of lung | 0 | 2 | 0.2051 | 0.0041 | 0.0117 | -0.0076 |
| pulmonary artery endothelial cell | 1 | 2 | 0.2118 | 0.0027 | 0.0018 | 0.0009 |
| mesothelial cell | 1 | 2 | 0.2276 | 0.0016 | 0.0006 | 0.0010 |
| non-classical monocyte | 1 | 2 | 0.2436 | 0.0055 | 0.0027 | 0.0028 |
| fibroblast of lung | 0 | 1 | 0.2465 | 0.0018 | 0.0026 | -0.0008 |
| neutrophil | 1 | 2 | 0.2657 | 0.0126 | 0.0093 | 0.0032 |
| pericyte | 0 | 1 | 0.2663 | 0.0001 | 0.0003 | -0.0001 |
| malignant cell | 0 | 2 | 0.3108 | 0.1160 | 0.1034 | 0.0126 |
| CD8-positive, alpha-beta T cell | 0 | 2 | 0.3560 | 0.0488 | 0.0747 | -0.0259 |
| conventional dendritic cell | 0 | 2 | 0.3692 | 0.0047 | 0.0061 | -0.0013 |
| classical monocyte | 1 | 2 | 0.4278 | 0.0769 | 0.0488 | 0.0281 |
| regulatory T cell | 0 | 2 | 0.4843 | 0.0176 | 0.0190 | -0.0014 |
| capillary endothelial cell | 1 | 2 | 0.4950 | 0.0023 | 0.0040 | -0.0017 |
| alveolar macrophage | 1 | 2 | 0.5328 | 0.1483 | 0.1382 | 0.0101 |
| plasmacytoid dendritic cell | 0 | 2 | 0.6519 | 0.0025 | 0.0048 | -0.0023 |
| pulmonary alveolar type 2 cell | 1 | 2 | 0.6719 | 0.0464 | 0.0236 | 0.0228 |
| mast cell | 0 | 1 | 0.8378 | 0.0072 | 0.0078 | -0.0060 |
| B cell | 1 | 2 | 0.8840 | 0.0904 | 0.0416 | 0.0488 |
| epithelial cell of lung | 1 | 2 | 0.9440 | 0.0549 | 0.0347 | 0.0202 |

Table 19: Posthoc Dunn Test of clustered LUAD patients at sub cell type level

#### 5.11 LUAD Clustering - Heatmap Visualization

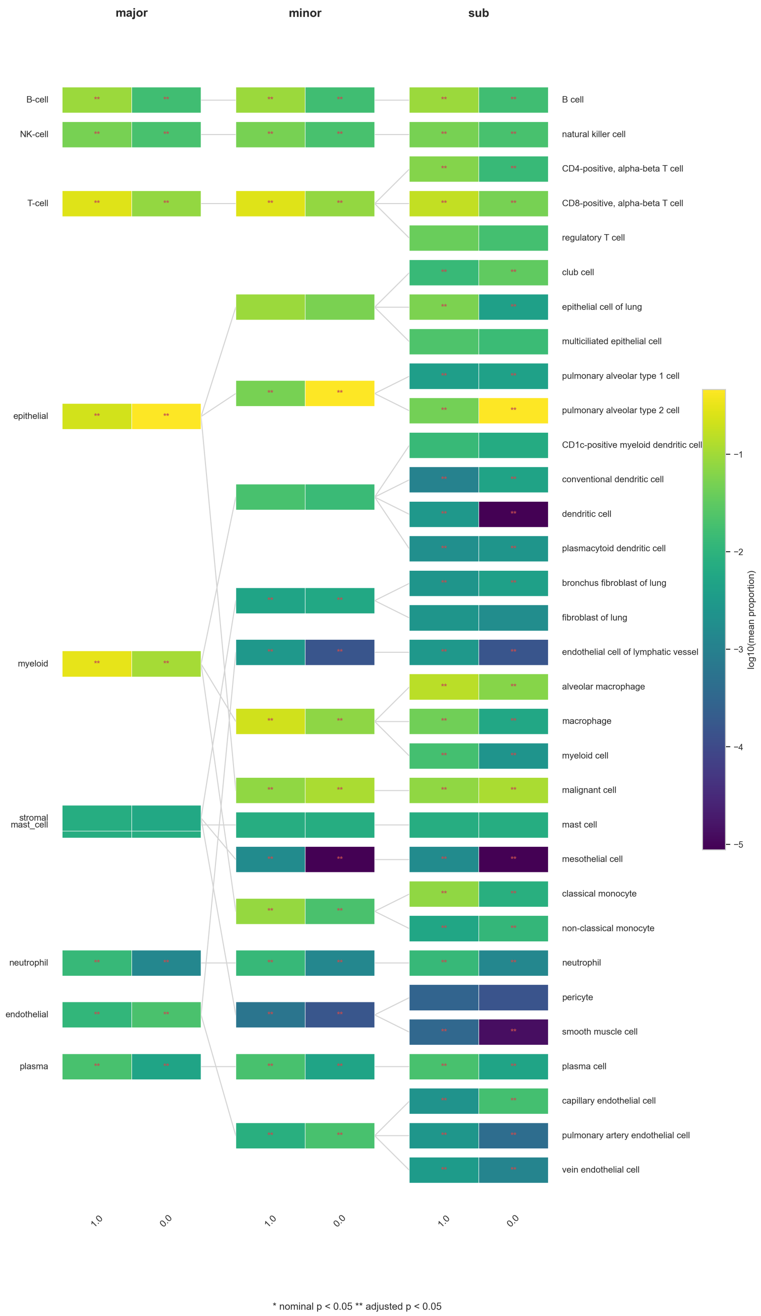

Figure 6: Heatmap visualization of log10 mean cellular proportions at all three cell hierarchy level for LUAD patient cluster 0 and cluster 1. Differences between cell types were evaluated with a Mann-Whitney-U test. Significance is highlighted by either "**" for adjusted p-values below 0.05 or "*" for nominal p-values below 0.05.

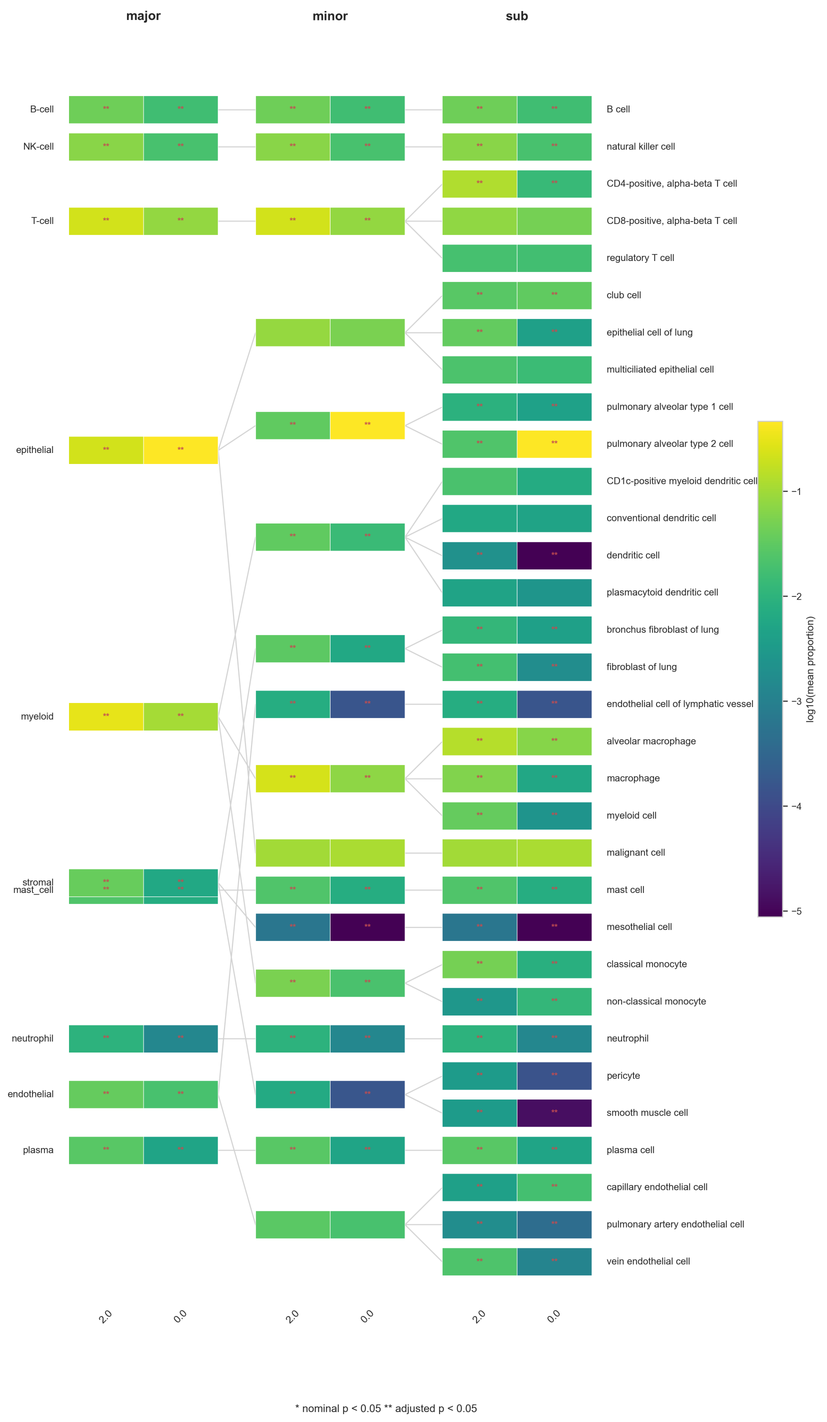

Figure 7: Heatmap visualization of log10 mean cellular proportions at all three cell hierarchy level for LUAD patient cluster 0 and cluster 2. Differences between cell types were evaluated with a Mann-Whitney-U test. Significance is highlighted by either "**" for adjusted p-values below 0.05 or "*" for nominal p-values below 0.05.

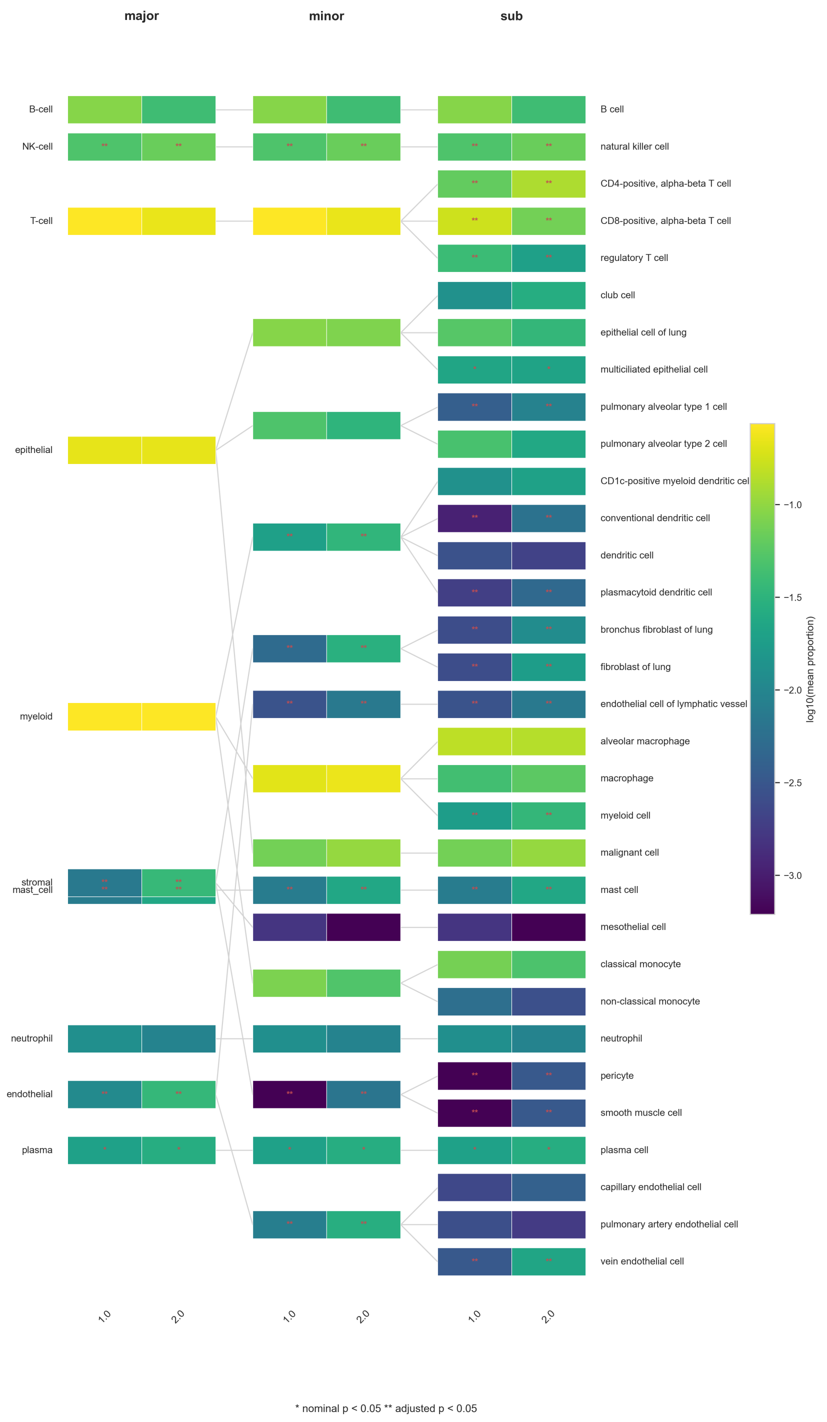

Figure 8: Heatmap visualization of log10 mean cellular proportions at all three cell hierarchy level for LUAD patient cluster 1 and cluster 2. Differences between cell types were evaluated with a Mann-Whitney-U test. Significance is highlighted by either "**" for adjusted p-values below 0.05 or "*" for nominal p-values below 0.05.

#### 5.12 LUAD Clustering - Hallmarks

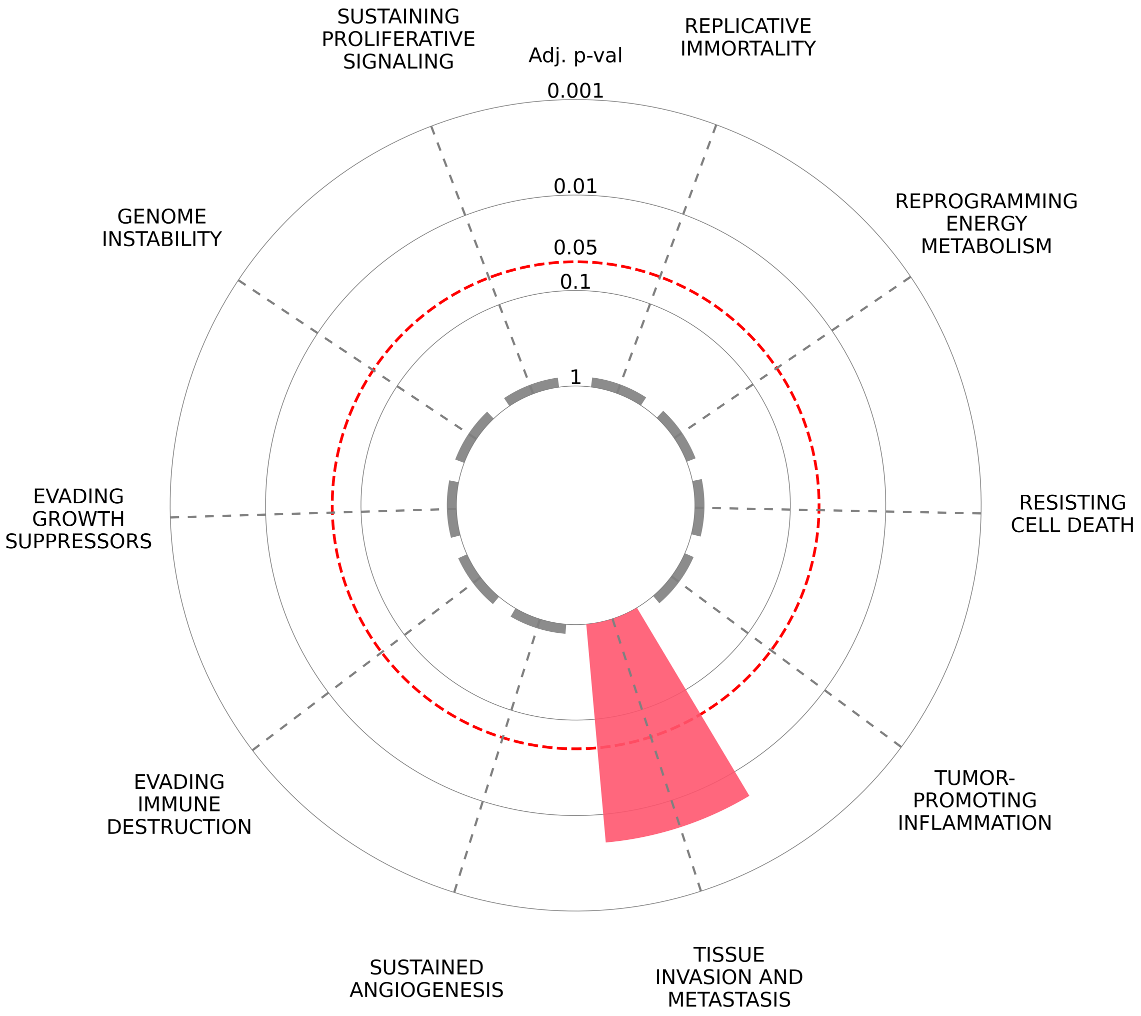

Figure 9: Cancer Hallmark enrichment analysis comparing the epithelial dominated TME state (cluster 0) and the stromal-immune TME state (cluster 2). The panel shows Hallmarks enriched in cluster 2 in comparison to cluster 0. Hallmarks located outside the red circle reached statistical significance.

#### 5.13 LUAD Clustering - Response Table

| | Cluster 0 | Cluster 1 | Cluster 2 |
|---|---|---|---|
| Non-response (PD) | 10 | 12 | 14 |
| Response (PR+CR) | 10 | 24 | 4 |

Table 20: Table showing number of patient respond state depending on the assigned cell composition cluster. Note that not all patients had information on their response state.

### 6 Sepsis Study

#### 6.1 Sepsis vs Healthy - Mann Whitney U Major

| celltype | sepsis mean ± std | healthy mean ± std | p | p_adj |
|---|---|---|---|---|
| B cell | $0.1129 \pm 0.0757$ | $0.0725 \pm 0.0308$ | 0.0039 | 0.0039 |
| DC | $0.0076 \pm 0.0074$ | $0.0302 \pm 0.0175$ | $1.29 \cdot 10^{-11}$ | $3.02 \cdot 10^{-11}$ |
| Monocyte | $0.3057 \pm 0.1327$ | $0.0877 \pm 0.0434$ | $6.90 \cdot 10^{-12}$ | $2.42 \cdot 10^{-11}$ |
| NK cell | $0.0500 \pm 0.0312$ | $0.1045 \pm 0.0643$ | $1.28 \cdot 10^{-5}$ | $1.49 \cdot 10^{-5}$ |
| Platelet | $0.0020 \pm 0.0026$ | $0.0001 \pm 0.0003$ | $2.26 \cdot 10^{-7}$ | $3.16 \cdot 10^{-7}$ |
| Progenitor cell | $0.0026 \pm 0.0018$ | $1.51 \cdot 10^{-5} \pm 6.48 \cdot 10^{-5}$ | $7.64 \cdot 10^{-16}$ | $5.35 \cdot 10^{-15}$ |
| T cell | $0.5193 \pm 0.1122$ | $0.7050 \pm 0.0758$ | $4.20 \cdot 10^{-11}$ | $7.35 \cdot 10^{-11}$ |

Table 21: Mann Whitney U test of sepsis vs healthy major cell type level

#### 6.2 Sepsis vs Healthy - Mann Whitney U Minor

| celltype | sepsis mean ± std | healthy mean ± std | p | p_adj |
|---|---|---|---|---|
| CD14 monocyte | $0.2429 \pm 0.1283$ | $0.0604 \pm 0.0431$ | $1.45 \cdot 10^{-11}$ | $6.04 \cdot 10^{-11}$ |
| CD16 monocyte | $0.0440 \pm 0.0420$ | $0.0078 \pm 0.0100$ | $1.57 \cdot 10^{-8}$ | $4.35 \cdot 10^{-8}$ |
| CD56bright NK cell | $0.0026 \pm 0.0034$ | $0.0046 \pm 0.0062$ | 0.4146 | 0.4319 |
| CD56dim NK cell | $0.0411 \pm 0.0312$ | $0.0993 \pm 0.0656$ | $4.90 \cdot 10^{-6}$ | $1.02 \cdot 10^{-5}$ |
| CD8aa | $0.0163 \pm 0.0143$ | $0.0064 \pm 0.0098$ | $7.39 \cdot 10^{-5}$ | 0.0001 |
| DN T cell | $0.0088 \pm 0.0140$ | $0.0020 \pm 0.0037$ | 0.0006 | 0.0009 |
| Effector B cell | $0.0186 \pm 0.0165$ | $0.0089 \pm 0.0156$ | 0.0008 | 0.0011 |
| Intermediate monocyte | $0.0187 \pm 0.0327$ | $0.0195 \pm 0.0211$ | 0.0811 | 0.0965 |
| MAIT | $0.0164 \pm 0.0216$ | $0.0537 \pm 0.0363$ | $5.00 \cdot 10^{-9}$ | $1.56 \cdot 10^{-8}$ |
| Memory B cell | $0.0192 \pm 0.03014$ | $0.0077 \pm 0.0089$ | 0.7069 | 0.7069 |
| Memory CD4 T cell | $0.1823 \pm 0.0850$ | $0.0664 \pm 0.0361$ | $5.26 \cdot 10^{-10}$ | $1.89 \cdot 10^{-9}$ |
| Memory CD8 T cell | $0.0343 \pm 0.0602$ | $0.1812 \pm 0.0930$ | $4.77 \cdot 10^{-13}$ | $3.97 \cdot 10^{-12}$ |
| Naive B cell | $0.0385 \pm 0.0296$ | $0.0515 \pm 0.0268$ | 0.0242 | 0.0303 |
| Naive CD4 T cell | $0.1214 \pm 0.1137$ | $0.3128 \pm 0.0899$ | $1.61 \cdot 10^{-12}$ | $1.00 \cdot 10^{-11}$ |
| Naive CD8 T cell | $0.0839 \pm 0.0437$ | $0.0403 \pm 0.0432$ | $4.19 \cdot 10^{-6}$ | $9.51 \cdot 10^{-6}$ |
| Plasma cell | $0.0172 \pm 0.0333$ | $0.0002 \pm 0.0005$ | $3.49 \cdot 10^{-12}$ | $1.75 \cdot 10^{-11}$ |
| Platelet | $0.0020 \pm 0.0026$ | $0.0001 \pm 0.0003$ | $2.26 \cdot 10^{-7}$ | $5.65 \cdot 10^{-7}$ |
| Progenitor cell | $0.0026 \pm 0.0018$ | $1.51 \cdot 10^{-5} \pm 6.48 \cdot 10^{-5}$ | $7.64 \cdot 10^{-16}$ | $1.91 \cdot 10^{-14}$ |
| Proliferating NK cell | $0.0063 \pm 0.0084$ | $0.0006 \pm 0.0010$ | $9.65 \cdot 10^{-5}$ | 0.0002 |
| Proliferating T cell | $0.0085 \pm 0.0157$ | $0.0023 \pm 0.0043$ | $1.35 \cdot 10^{-5}$ | $2.60 \cdot 10^{-5}$ |
| Transitional B cell | $0.0193 \pm 0.0362$ | $0.0042 \pm 0.0069$ | 0.0002 | 0.0002 |
| Treg | $0.0154 \pm 0.0144$ | $0.0104 \pm 0.0069$ | 0.1834 | 0.2084 |
| cDC2 | $0.0045 \pm 0.0067$ | $0.0293 \pm 0.0174$ | $3.35 \cdot 10^{-13}$ | $3.97 \cdot 10^{-12}$ |
| gdT | $0.0320 \pm 0.0254$ | $0.0296 \pm 0.0326$ | 0.262 3 | 0.2851 |
| pDC | $0.0030 \pm 0.0041$ | $0.0009 \pm 0.0009$ | 0.0023 | 0.0031 |

Table 22: Mann Whitney U test of sepsis vs healthy minor cell type level

#### 6.3 Sepsis vs Healthy - Heatmap Visualization

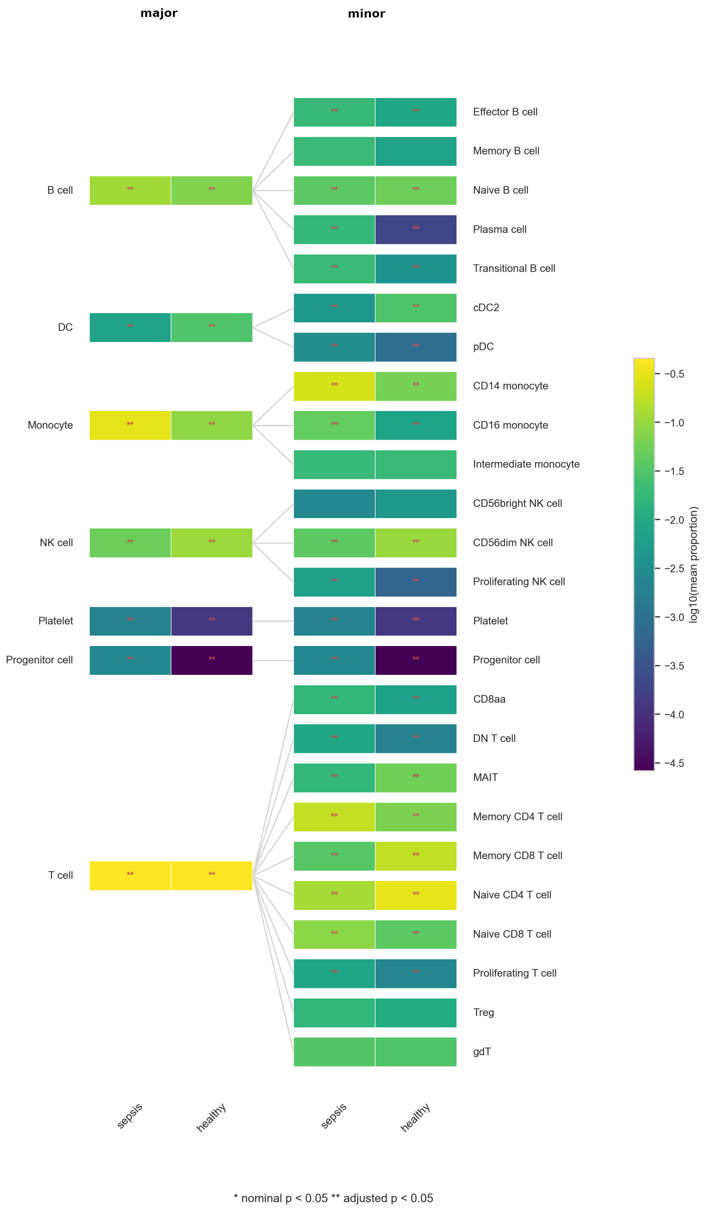

Figure 10: Heatmap visualization of log10 mean cellular proportions for the sepsis and healthy PBMC cohort. Differences between cell types were evaluated with a Mann-Whitney-U test. Significance is highlighted by either "**" for adjusted p-values below 0.05 or "*" for nominal p-values below 0.05.

### 7 COVID19 Study

#### 7.1 Covid vs Healthy - Mann Whitney U Major

| celltype | control<br>mean ± std | covid<br>mean ± std | p | p_adj |
|---|---|---|---|---|
| B cell | $0.0995 \pm 0.0316$ | $0.1060 \pm 0.0447$ | 0.8904 | 0.8904 |
| DC | $0.0072 \pm 0.0066$ | $0.0077 \pm 0.0051$ | 0.5815 | 0.8142 |
| Monocyte | $0.1645 \pm 0.0512$ | $0.2083 \pm 0.1048$ | 0.4909 | 0.8142 |
| NK cell | $0.0780 \pm 0.0393$ | $0.0800 \pm 0.0323$ | 0.7048 | 0.8222 |
| Platelet | $2.00 \cdot 10^{-5} \pm 8.24 \cdot 10^{-5}$ | $0.0020 \pm 0.0029$ | 0.0003 | 0.0011 |
| Progenitor cell | $0.0 \pm 0.0$ | $0.0021 \pm 0.0021$ | $4.82 \cdot 10^{-6}$ | $3.37 \cdot 10^{-5}$ |
| T cell | $0.6508 \pm 0.0690$ | $0.5939 \pm 0.1258$ | 0.1296 | 0.3025 |

Table 23: Mann Whitney U test of covid vs healthy major cell type level

#### 7.2 Covid vs Healthy - Mann Whitney U Minor

| celltype | control<br>mean ± std | covid<br>mean ± std | p | p_adj |
|---|---|---|---|---|
| CD14 monocyte | $0.1387 \pm 0.0544$ | $0.1714 \pm 0.1106$ | 0.6297 | 0.6844 |
| CD16 monocyte | $0.0156 \pm 0.0153$ | $0.0229 \pm 0.0193$ | 0.3348 | 0.4406 |
| CD56bright NK cell | $0.0042 \pm 0.0037$ | $0.0071 \pm 0.0052$ | 0.1385 | 0.2164 |
| CD56dim NK cell | $0.0735 \pm 0.0398$ | $0.0615 \pm 0.0329$ | 0.4486 | 0.5607 |
| CD8aa | $0.0030 \pm 0.0058$ | $0.0066 \pm 0.0081$ | 0.2229 | 0.3096 |
| DN T cell | $0.0031 \pm 0.0065$ | $0.0152 \pm 0.0125$ | 0.0009 | 0.0023 |
| Effector B cell | $0.0025 \pm 0.0057$ | $0.0170 \pm 0.0210$ | 0.0025 | 0.0058 |
| Intermediate monocyte | $0.0101 \pm 0.0117$ | $0.0140 \pm 0.0226$ | 0.6046 | 0.6844 |
| MAIT | $0.0418 \pm 0.0278$ | $0.0046 \pm 0.0067$ | $1.19 \cdot 10^{-5}$ | $5.93 \cdot 10^{-5}$ |
| Memory B cell | $0.0258 \pm 0.0190$ | $0.0123 \pm 0.0150$ | 0.0157 | 0.0303 |
| Memory CD4 T cell | $0.1124 \pm 0.0525$ | $0.2488 \pm 0.0532$ | $4.63 \cdot 10^{-6}$ | $3.39 \cdot 10^{-5}$ |
| Memory CD8 T cell | $0.0666 \pm 0.0768$ | $0.0668 \pm 0.0784$ | 0.7559 | 0.7874 |
| Naive B cell | $0.0655 \pm 0.0227$ | $0.0315 \pm 0.0267$ | 0.0003 | 0.0010 |
| Naive CD4 T cell | $0.2656 \pm 0.0654$ | $0.1168 \pm 0.1068$ | 0.0004 | 0.0012 |
| Naive CD8 T cell | $0.0729 \pm 0.0500$ | $0.0681 \pm 0.0304$ | 0.9725 | 0.9725 |
| Plasma cell | $0.0 \pm 0.0$ | $0.0266 \pm 0.0239$ | $4.27 \cdot 10^{-7}$ | $1.07 \cdot 10^{-5}$ |
| Platelet | $2.00 \cdot 10^{-5} \pm 8.24 \cdot 10^{-5}$ | $0.0020 \pm 0.0029$ | 0.0003 | 0.0010 |
| Progenitor cell | $0.0 \pm 0.0$ | $0.0021 \pm 0.0021$ | $4.82 \cdot 10^{-6}$ | $3.39 \cdot 10^{-5}$ |
| Proliferating NK cell | $0.0002 \pm 0.0006$ | $0.0114 \pm 0.0123$ | $3.74 \cdot 10^{-5}$ | 0.0002 |
| Proliferating T cell | $0.0051 \pm 0.0063$ | $0.0167 \pm 0.0161$ | 0.0130 | 0.0271 |
| Transitional B cell | $0.0058 \pm 0.0062$ | $0.0185 \pm 0.0174$ | 0.0475 | 0.0849 |
| Treg | $0.0189 \pm 0.0161$ | $0.0148 \pm 0.0175$ | 0.1902 | 0.2798 |
| cDC2 | $0.0064 \pm 0.0065$ | $0.0025 \pm 0.0036$ | 0.0527 | 0.0879 |
| gdT | $0.0615 \pm 0.0700$ | $0.0355 \pm 0.0246$ | 0.5128 | 0.6105 |
| pDC | $0.0007 \pm 0.0008$ | $0.0052 \pm 0.0036$ | $5.42 \cdot 10^{-6}$ | $3.9 \cdot 10^{-5}$ |

Table 24: Mann Whitney U test of covid vs healthy sub cell type level

##### 7.3 Covid vs Healthy - Heatmap Visualization

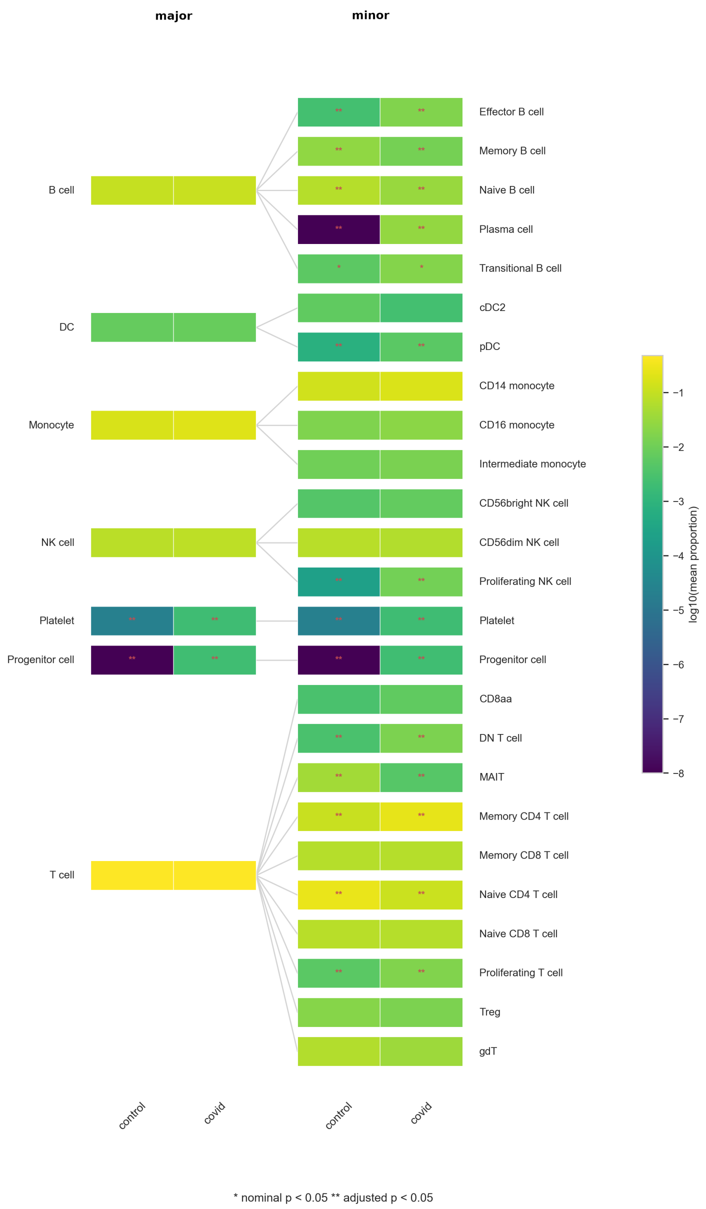

Figure 11: Heatmap visualization of log10 mean cellular proportions for the covid and healthy PBMC cohort. Differences between cell types were evaluated with a Mann-Whitney-U test. Significance is highlighted by either "**" for adjusted p-values below 0.05 or "*" for nominal p-values below 0.05.

### 8 Lupus Study

#### 8.1 Lupus vs Healthy - Mann Whitney U Major

| celltype | Lupus mean ± std | Healthy mean ± std | p | p_adj |
|---|---|---|---|---|
| B cell | $0.1013 \pm 0.0450$ | $0.0890 \pm 0.0303$ | 0.2486 | 0.2900 |
| DC | $0.0121 \pm 0.0109$ | $0.0105 \pm 0.0088$ | 0.7673 | 0.7673 |
| Monocyte | $0.2319 \pm 0.0871$ | $0.1494 \pm 0.0563$ | 0.0002 | 0.0011 |
| NK cell | $0.0718 \pm 0.0462$ | $0.0847 \pm 0.0361$ | 0.1313 | 0.2298 |
| Platelet | $0.0007 \pm 0.0014$ | $0.0006 \pm 0.0007$ | 0.1068 | 0.2298 |
| Progenitor cell | $0.0003 \pm 0.0009$ | $8.32 \cdot 10^{-5} \pm 0.0003$ | 0.1731 | 0.2423 |
| T cell | $0.5819 \pm 0.1097$ | $0.6657 \pm 0.0816$ | 0.00309 | 0.0108 |

Table 25: Mann Whitney U test of lupus vs healthy major cell type level

#### 8.2 Lupus vs Healthy - Mann Whitney U Minor

| celltype | Lupus mean ± std | Healthy mean ± std | p | p_adj |
|---|---|---|---|---|
| CD14 monocyte | $0.1833 \pm 0.0783$ | $0.1006 \pm 0.0626$ | $9.91 \cdot 10^{-5}$ | 0.0002 |
| CD16 monocyte | $0.0377 \pm 0.0199$ | $0.0133 \pm 0.0140$ | $5.56 \cdot 10^{-6}$ | $2.78 \cdot 10^{-5}$ |
| CD56bright NK cell | $0.0102 \pm 0.0090$ | $0.0045 \pm 0.0064$ | 0.0022 | 0.0046 |
| CD56dim NK cell | $0.0554 \pm 0.0482$ | $0.0762 \pm 0.0373$ | 0.0563 | 0.0988 |
| CD8aa | $0.0058 \pm 0.0076$ | $0.0063 \pm 0.0100$ | 0.5389 | 0.5914 |
| DN T cell | $0.0166 \pm 0.0126$ | $0.0026 \pm 0.0047$ | $2.12 \cdot 10^{-7}$ | $1.77 \cdot 10^{-6}$ |
| Effector B cell | $0.0153 \pm 0.0146$ | $0.0160 \pm 0.0120$ | 0.5441 | 0.5914 |
| Intermediate monocyte | $0.0110 \pm 0.0195$ | $0.0355 \pm 0.0276$ | 0.0004 | 0.0010 |
| MAIT | $0.0085 \pm 0.0134$ | $0.0560 \pm 0.0515$ | $8.53 \cdot 10^{-6}$ | $3.05 \cdot 10^{-5}$ |
| Memory B cell | $0.0103 \pm 0.0132$ | $0.0154 \pm 0.0143$ | 0.0593 | 0.0988 |
| Memory CD4 T cell | $0.2084 \pm 0.0592$ | $0.0903 \pm 0.0630$ | $4.80 \cdot 10^{-8}$ | $1.07 \cdot 10^{-6}$ |
| Memory CD8 T cell | $0.0852 \pm 0.0587$ | $0.0734 \pm 0.0784$ | 0.1829 | 0.2406 |
| Naive B cell | $0.0675 \pm 0.0316$ | $0.0423 \pm 0.0271$ | 0.0016 | 0.0040 |
| Naive CD4 T cell | $0.1476 \pm 0.1125$ | $0.2609 \pm 0.0691$ | $7.92 \cdot 10^{-6}$ | $3.05 \cdot 10^{-5}$ |
| Naive CD8 T cell | $0.0541 \pm 0.0453$ | $0.0881 \pm 0.0490$ | 0.0088 | 0.0170 |
| Plasma cell | $0.0025 \pm 0.0042$ | $0.0010 \pm 0.0023$ | 0.3393 | 0.4241 |
| Platelet | $0.0007 \pm 0.0014$ | $0.0006 \pm 0.0007$ | 0.1068 | 0.1571 |
| Progenitor cell | $0.0003 \pm 0.0009$ | $8.32 \cdot 10^{-5} \pm 0.0003$ | 0.1731 | 0.2404 |
| Proliferating NK cell | $0.0062 \pm 0.0132$ | $0.0041 \pm 0.0057$ | 0.3864 | 0.4600 |
| Proliferating T cell | $0.0104 \pm 0.0166$ | $0.0062 \pm 0.0148$ | 0.0677 | 0.1057 |
| Transitional B cell | $0.0057 \pm 0.0138$ | $0.0142 \pm 0.0106$ | $2.15 \cdot 10^{-6}$ | $1.34 \cdot 10^{-5}$ |
| Treg | $0.0291 \pm 0.0298$ | $0.0111 \pm 0.0125$ | 0.0017 | 0.0040 |
| cDC2 | $0.0100 \pm 0.0100$ | $0.0084 \pm 0.0075$ | 0.7329 | 0.7634 |
| gdT | $0.0163 \pm 0.0133$ | $0.0707 \pm 0.0562$ | $8.56 \cdot 10^{-8}$ | $1.07 \cdot 10^{-6}$ |
| pDC | $0.0021 \pm 0.0018$ | $0.0021 \pm 0.0019$ | 0.9056 | 0.9056 |

Table 26: Mann Whitney U test of lupus vs healthy sub cell type level

#### 8.3 Lupus vs Healthy - Heatmap Visualization

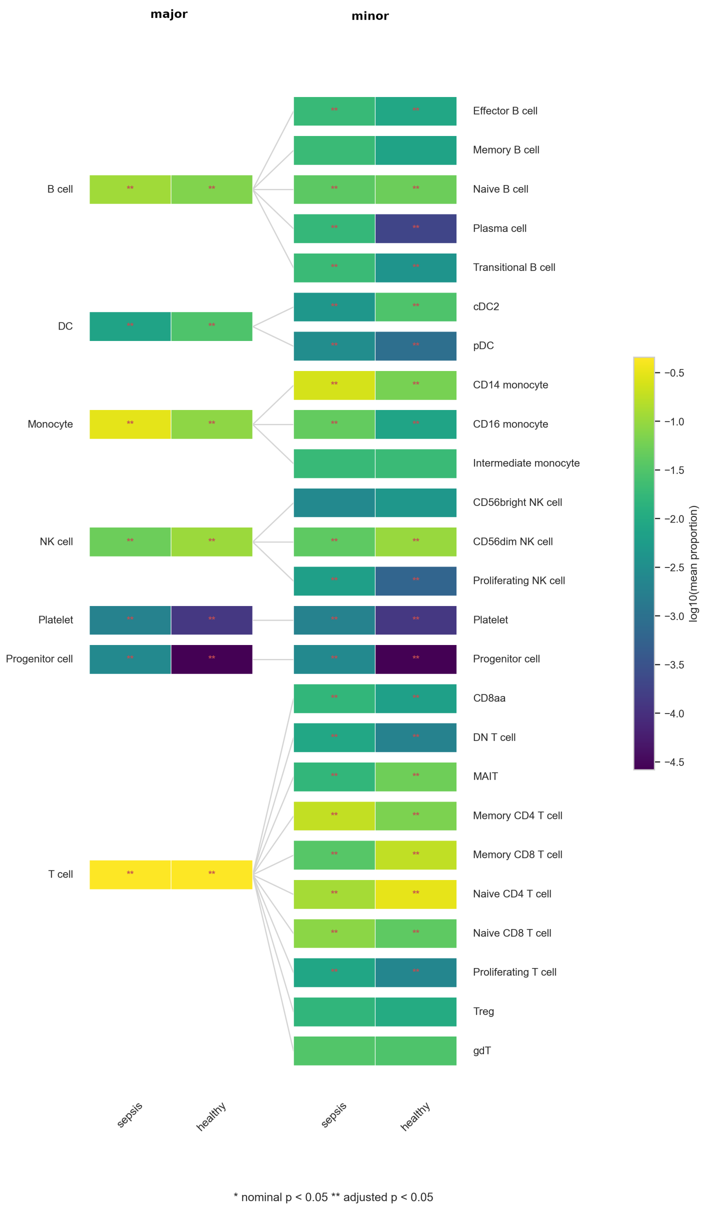

Figure 12: Heatmap visualization of log10 mean cellular proportions for the lupus and healthy PBMC cohort. Differences between cell types were evaluated with a Mann-Whitney-U test. Significance is highlighted by either "**" for adjusted p-values below 0.05 or "*" for nominal p-values below 0.05.
